# Individual behavioral trajectories shape whole-brain connectivity in mice

**DOI:** 10.1101/2022.03.25.485806

**Authors:** Jadna Bogado Lopes, Anna N. Senko, Klaas Bahnsen, Daniel Geisler, Eugene Kim, Michel Bernanos, Diana Cash, Stefan Ehrlich, Anthony C. Vernon, Gerd Kempermann

## Abstract

It is widely assumed that our actions shape our brains and that the resulting connections determine who we are. To test this idea in a reductionist setting, in which genes and environment are controlled, we investigated differences in neuroanatomy and structural covariance by *ex vivo* structural magnetic resonance imaging (MRI) in mice whose behavioral activity was continuously tracked for 3 months in a large, enriched environment. We confirmed that environmental enrichment increases mouse hippocampal volumes. Stratifying the enriched group according to individual longitudinal behavioral trajectories, however, revealed striking differences in mouse brain structural covariance in continuously highly active mice compared to those whose trajectories showed signs of habituating activity. Network-based statistics identified distinct sub-networks of murine structural covariance underlying these differences in behavioral activity. Together, these results reveal that differentiated behavioral trajectories of mice in an enriched environment are associated with differences in brain connectivity.

## Introduction

The ‘non-shared environment’ (1), the elusive component of the non-genetic factor in phenotypic variation (2), contributes to inter-individual differences and neurobiological individuality. In humans, *in-vivo* neuroimaging studies have demonstrated a complex relationship between behavioral performance and brain structure associated with learning (3, 4), personality traits (5), and political orientation (6), supporting the idea of a causal, yet individual relationship between brain function, structure and behavior. The structural connectome, which can be estimated based on the correlation between regional brain volumes (across subjects) or white matter links between specific brain regions (within subjects) partially recapitulates known functional networks and represents an individual ‘fingerprint’ in humans and non-human animals (7–10). We have reported that environmental enrichment (ENR) increases variability in activity-related behaviors (roaming entropy, RE), or total object exploration in a novel object recognition task (11). The ENR mice also showed greater variability in measures linked to brain plasticity, such as adult hippocampal neurogenesis (11) accompanied by distinguishing epigenetic patterns in the hippocampal dentate gyrus (12, 13).

Structural magnetic resonance imaging (MRI) studies of mice exposed to different enrichment paradigms have confirmed historical *post-mortem* findings, such as increased hippocampal volumes (13), and extended these to mouse brain regions involved in sensorimotor processing (14). We here go a decisive step further and examine the relationships between individual behavioral trajectories and whole brain structural *networks* (“connectomics”). These data may provide neurobiological foundations for concepts such as cognitive reserve or brain maintenance, which attempt to capture individual differences in healthy cognitive aging and resilience to neurodegenerative disease (15, 16).

## Results

We longitudinally tracked the behavior of enriched female mice (n = 38) in our Individuality cage system, equipped with radio-frequency identification (RFID) technology (12) (Fig. 1A). During 12-weeks of exposure to the ENR, mean values of roaming entropy (RE) as a measure of exploration and territorial coverage (12, 17) were calculated per night and aggregated across four time-blocks of 21 nights each. Increasing interindividual component of variance (Fig. 1B) confirmed the emergence of individual behavioral differences. Two distinguishable patterns were observed (Fig. 1C): mice with consistently high levels of RE and those with habituation and decreased RE values over time. Mice were stratified according to this criterion and grouped into ‘flat’ roamers (n = 15) or ‘down’ roamers (n = 15) depending on the slope of the linear regression line through a set of four RE time-blocks. The slopes for ‘flat’ roamers ranged from -0.003 to 0.004, while slopes lower than -0.006 (to -0.049) were considered to represent ‘down’ roamers. Mice that showed in-between slope values (n = 10) were excluded from the clustering to achieve a sharper distinction.

**Figure 1.**
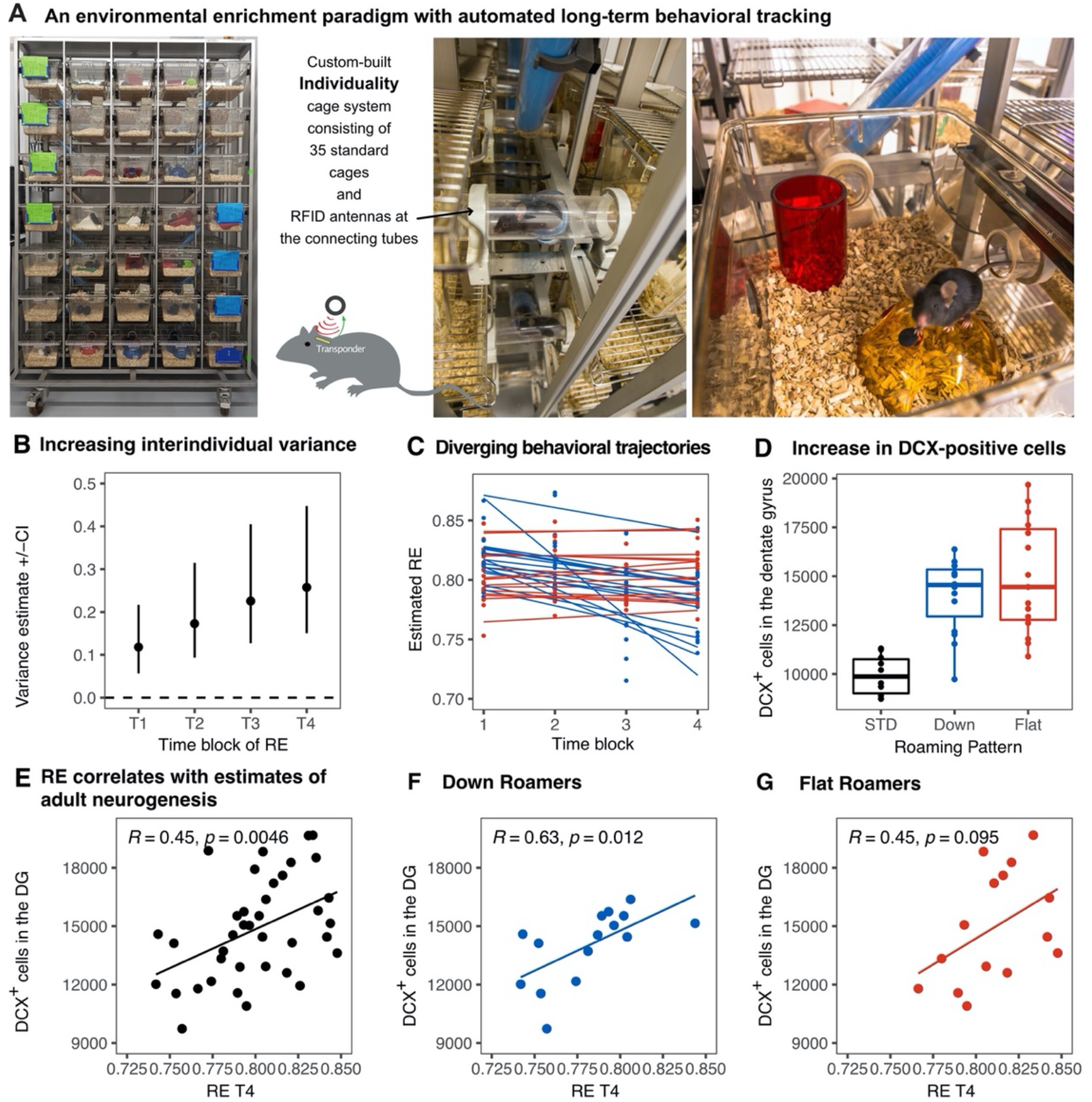
Emergence of inter-individual differences after environmental enrichment. Pictures **(A)** of the “individuality” cage—a custom build RFID cage with 70 interconnected small cages and 115 antennae distributed in the connecting tunnels. Interindividual variance **(B)** of behavior increased over time. Subgrouping of mice (n = 15) based on the slopes of RE trajectories **(C)**, blue: ‘down’, and red: ‘flat’. Animals living under enriched environment conditions (ENR) showed increased adult hippocampal neurogenesis (as assessed by proxy marker DCX) independent of behavioral trajectories (One-way ANOVA: F_(2,37)_ = 17.3, p < 0.001; Tukey post-hoc: standard vs flat and standard vs down p < 0.01, flat vs down p = 0.46) **(D)**. Correlations between the number of doublecortin positive cells in the dentate gyrus to RE values at time-block four in the enriched group (R^2^ = 0.20, p = 0.005) **(E)**, down subgroup (R^2^ = 0.39, p = 0.012) **(F)**, and flat subgroup (R^2^ = 0.20, p = 0.095) **(G)**. Box and whisker plots: center line - median; upper and lower hinges - first and third quartiles; whiskers - highest and lowest values within 1.5 times the interquartile range outside hinges; dots - individual data points.

The number of doublecortin (DCX) positive cells in the dentate gyrus, a proxy measure of adult neurogenesis, increased in ENR compared to standard conditions, but did not differ between the ‘flat’ vs. ‘down’ ENR subgroups (Fig. 1D). As previously shown (12, 17), individual variability in behavior positively correlated to adult hippocampal neurogenesis (R^2^ = 0.20, p = 0.0046; Fig. 1E). The positive correlation between end-point behavior (RE at the time-block four) and immature neurons was statistically significant in the ‘down’ roamers subgroup (R^2^ = 0.40, p = 0.012), but missed conventional statistical significance in the ‘flat’ roamers (R^2^ = 0.20, p = 0.095; Fig. 1F-G). However, the correlations in the flat and down subgroups were not statistically different (one-sided Fisher’s *z* test, *z* = - 0.63, *p* = 0.27), which might point to an insufficient power for this comparison.

We next explored neuroanatomical changes between ENR and standard-housed (STD) mice by *ex vivo* structural MRI. Total brain volumes did not differ significantly (STD: 450.3±11.7 mm^3^ vs. ENR: 456.5±16.0 mm^3^; *t* = 1.14, *df* = 46; p > 0.05; q > 0.05; Fig. S1A). Using atlas-based segmentation, statistically significant (5 % False Discovery Rate [FDR]) group differences in absolute volume (mm^3^) were found for 12 % (22/182) of mouse brain atlas regions of interest (ROI) with effect sizes (SMD) ranging from +2.7 in CA1 oriens (CA1Or) to +1.15 in the molecular layer of the dentate gryus (MoDG; Fig. 2a; Table S1). Most of these ROIs (68 %; 15/22) were located in the mouse hippocampus, but absolute volume increases were also observed in the infralimbic cortex (cingulate cortex area 25), olfactory nuclei, the anterior commissure and the orbital cortex (Fig. 2a; Table S1). The majority of these ROIs were conserved as significant group differences when considering relative volumes (% of whole brain), suggesting normal scaling (Fig. S2A; Table S2). Complementing the atlas-based analysis, voxel-wise assessment of volume changes using tensor based morphometry (TBM) revealed statistically significant (Family Wise Error rate p<0.05) clusters of voxels with apparent increases in both absolute volumes (Fig. S2B) and relative volumes (Fig. S2C) localized within the mouse hippocampus when comparing ENR to STD mice. Collectively, these data confirm the expected effects of ENR on mouse brain structure, as evidenced by prior mouse neuroimaging (13, 14) and historical *post-mortem* studies in rats (18).

**Figure 2.**
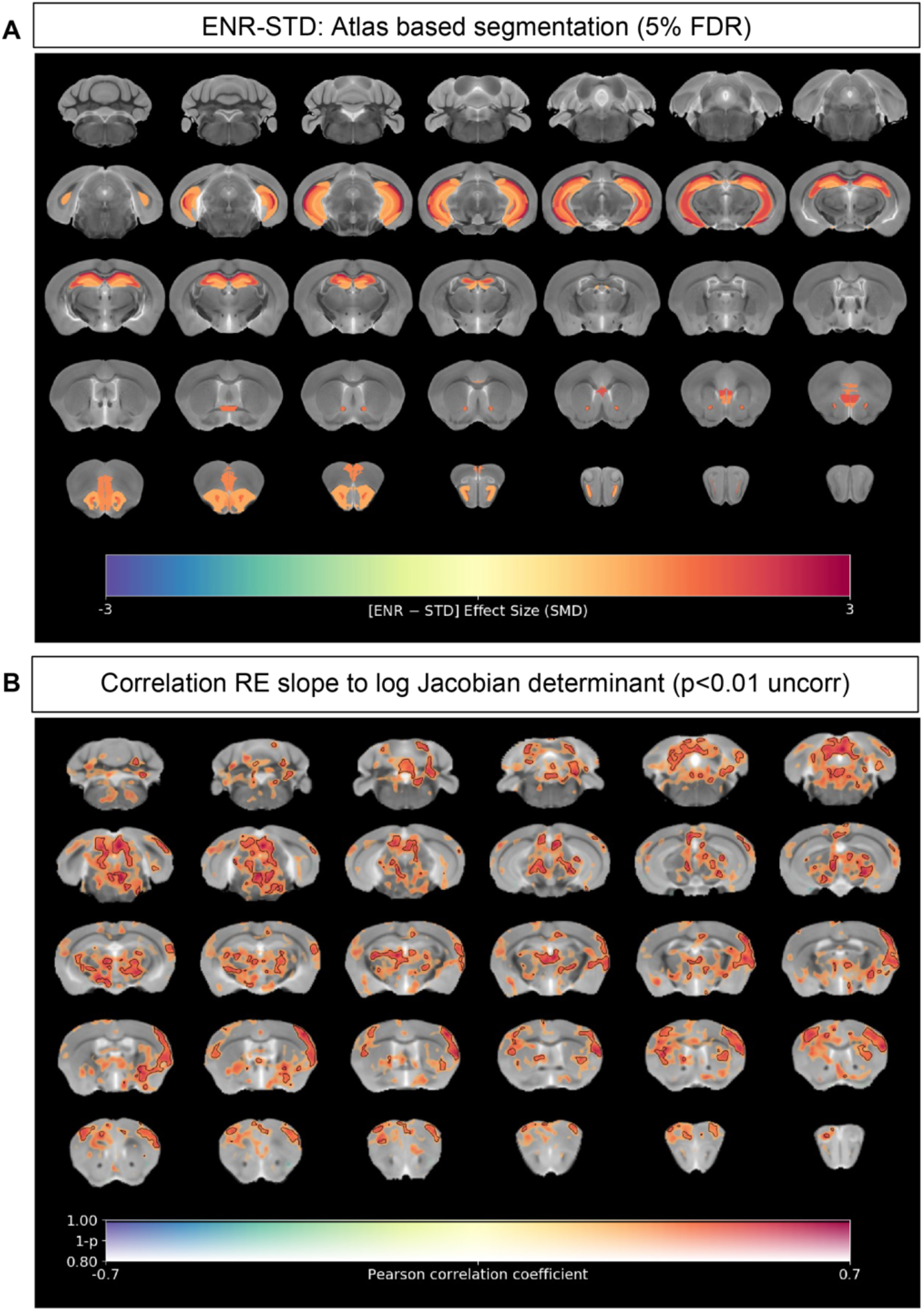
Neuroanatomical differences associated with exposure to an enriched environment and with variation in roaming entropy. **(A)** Map of regional differences in absolute volume (mm3) extracted using the DSURQE mouse brain atlas comparing ENR mice to those in standard (STD) housing. Only atlas regions of interest (ROI) that survive multiple comparisons correction are shown (False Discovery Rate, 5%). Data shown for each statistically significant ROI is the standardized mean difference (SMD) comparing ENR – STD housed mice. Warm colors indicate regional volume increases, whilst cold colors indicate regional volume decreases in ENR-mice as compared to those in STD housing. **(B)** Positive correlation (Pearson’s *r*) between roaming entropy slope across all four time blocks and local brain absolute volume changes (log-scaled Jacobian determinant) for mice clustering into both ‘flat’ and ‘down’ groups. Voxel clusters with solid contours are significantly correlated to behavior at an exploratory threshold of uncorrected p < 0.01. No changes were detected after correction for multiple comparisons (FWE p < 0.05). Warm colors indicate positive correlation, suggesting increased volumes in ‘flat’ as compared to ‘down’ mice.

Total brain volumes did not differ between the two ENR sub-groups (‘down’: 451.7±11.8 mm^3^ vs. ‘flat’: 459.2±5.2 mm^3^; *t* = 1.24, *df* = 28; p > 0.05; q > 0.05; Fig. S1B). Voxel-wise correlation of the slope of the linear regression line across the four RE time-blocks against log-Jacobian measures of local volume change for each individual mouse in the ‘flat’ and ‘down’ sub-groups revealed that RE slope values correlated positively with the absolute volumes of cingulate, motor, somatosensory, insular, visual and auditory cortices, but also the ventral thalamic nuclei and several mid- and hind-brain regions, at a threshold of p < 0.01 uncorrected for multiple comparisons (Fig. 2B). These data suggest that in ‘down’ roamers, with negative RE slope values, the absolute volumes of these regions were smaller than in the mice that displayed a ‘flat’ RE behavior. Exploratory voxel-wise analysis of local volume changes between ‘flat’ and ‘down’ roamers (Fig. S3) similarly highlighted clusters of voxels with apparent increases in volume in the ‘flat’ relative to the ‘down’ sub-group in the cingulate, motor, somatosensory, insular, visual and auditory cortices, ventral thalamic nuclei and several mid- and hind-brain regions, largely replicating the pattern in Fig. 3B. None of these clusters remained statistically significant after correction for multiple comparisons (FWE p < 0.05). Hence, whilst these data will require replication in a larger cohort, they corroborate the positive association between RE slope and brain volume, suggesting that ‘flat’ roamers may have larger volumes of these brain structures as compared to ‘down’ roamers. Collectively, these data also suggest that individual differences in RE behaviors may be supported by widely distributed mouse brain circuitry.

**Figure 3.**
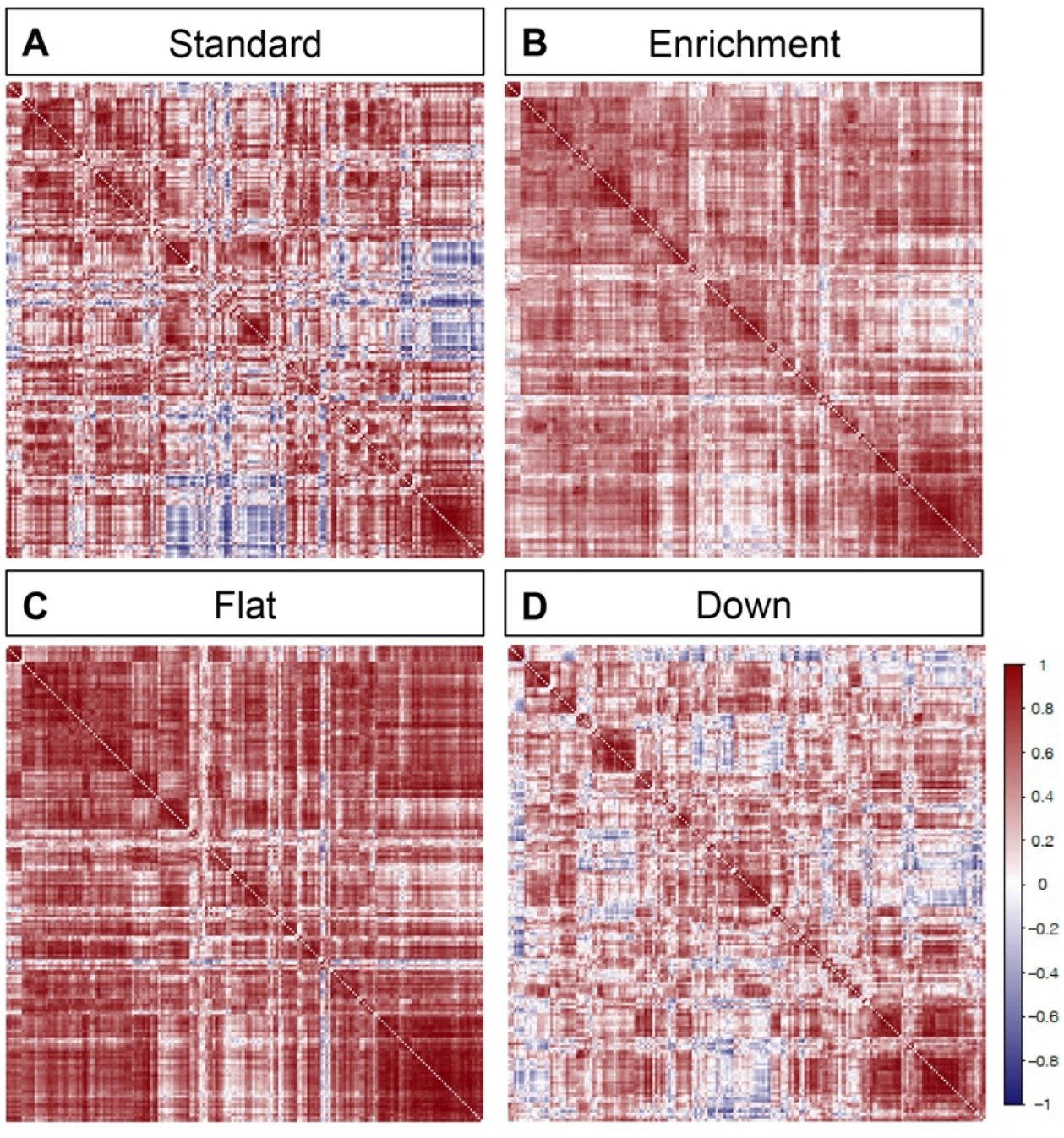
Environmental enrichment distinctively alters brain structural covariance. Rows and columns of each matrix denote atlas-defined structures, and color intensity the correlation strength (Pearson’s correlation, red = positive, blue = negative). Mice were kept either in **(A)** STD or **(B)** ENR conditions. Furthermore, based on patterns of behavior, ENR animals were subdivided into **(C)** ‘flat’ and **(D)** ‘down’ roamers. Labels indicate housing (top panels) or ENR subgroups (bottom panels).

To study whether the emergence of inter-individual variation in RE was associated with increased variability in the volumes of mouse brain regions, we calculated the coefficient of variation (CV) for each individual brain region. To determine if there are different degrees of overall variability between ENR subgroups, we averaged the CVs across all 182 ROIs in the mouse brain atlas to yield an average variability measure for each group (Fig. S4). Comparing the sum of ranks between ‘down’ and ‘flat’ mice, we found a highly statistically significant difference (Mann-Whitney U = 11076; p < 0.0001; Fig. S4). These data are indicative of an increase in regional brain volume variance as a function of behavioral response to the enriched environment.

Structural covariance is defined as the correlated variation in volumes between pairs of brain regions, which, as human neuroimaging studies suggest, reflects both structural (19) and functional brain connectivity (20). Structural covariance networks are conserved in the mouse brain, providing an opportunity to explore the impact of ENR compared to standard housing on this cross-species measure of brain connectivity (9, 10). Correlation matrices of mouse brain structural covariance (Fig. 3A-B) revealed a broad effect of housing, which was found to be statistically significant (Chi square tests for equality of two correlation matrices; Chi Square = 17997.7, df = 16471, *p* < 0.0001, where *prob* is the probability of observing the Chi Square under the null hypothesis) (21). Furthermore, comparison of structural covariance matrices within the ENR group revealed distinctive effects between ‘down’ and ‘flat’ roamers (Chi Square = 19997.29, df = 16471, p < 0.0001; Fig. 3C-D). The structural covariance matrix of ‘flat’ roamers (Fig. 3C) differed from that of STD animals (Chi Square = 17938.04, df = 16471, prob < 0.0001), whereas ‘down’ roamers (Fig. 3D) were highly similar to STD group (Chi Square = 14327.89, df = 16471, prob < 1) (Fig. 3A). In general, brain regions in the ‘flat’ roamers were highly correlated with each other, whereas this was not the case in ‘down’ roamers and STD mice (Fig. 3). These correlation patterns support the hypothesis of a link between behavioral trajectories and distinct levels of brain structural covariance, which develops independently of genetic variation, or the idea that emerging complex behavioral patterns are dependent on changes in covariance between widely distributed regions.

To test hypotheses regarding specific network connections we used Network based statistics (NBS). To describe the graph model, an appropriate set of nodes was defined using the mouse brain ROIs from the DSURQE atlas that differed significantly in volumes based on the ENR>Standard contrast after correction for multiple comparisons (5 % FDR; Fig. 2A, Table S1). Values of bilateral regions were averaged and plotted in both hemispheres. Comparing ENR and STD mice, NBS analysis detected a single statistically significant sub-network of increased structural covariance, comprising 23 structural connections (*t* = 2.4; *p* < 0.05). By contrast, comparing ‘flat’ vs. ‘down’ mice in the ENR group, NBS identified a larger statistically significant sub-network of increased structural covariance comprising 49 structural connections (*t* = 2.4; *p* < 0.05; Fig. 4A, B and Table S3). The NBS analysis supports the conclusion that inter-individual differences in behavioral trajectories (here clustered into the ‘flat’ vs. ‘down’ roamer subgroups of ENR) surpassed the housing effects, and identifies specific sub-networks of structural covariance in the mouse brain.

**Figure 4.**
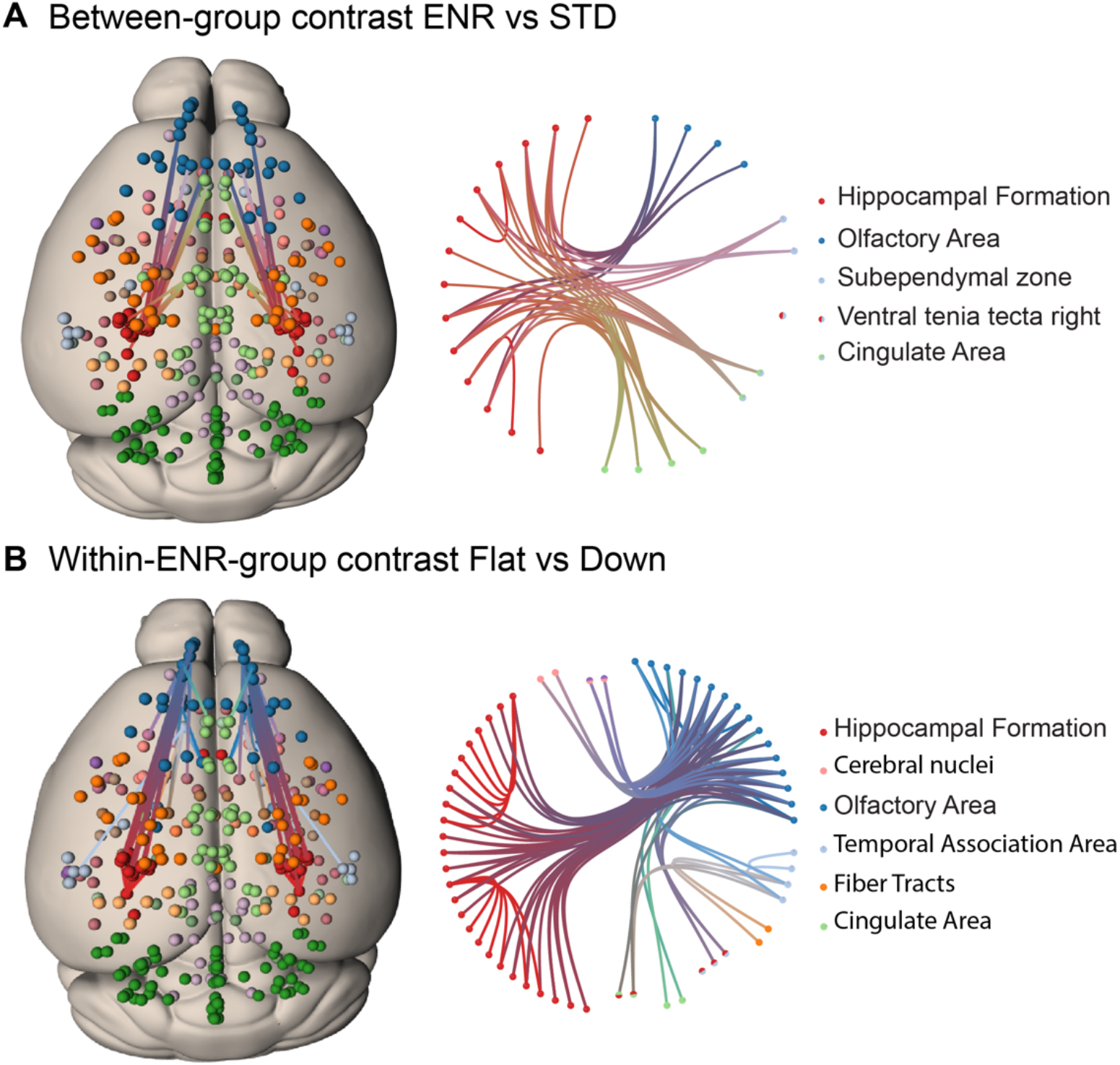
Network based statistics confirms differential structural connectivity patterns. Visualization of subnetworks in the contrasts **(A)** ENR > STD and **(B)** FLAT > DOWN both at a threshold of t = 2.4. In the left panels nodes are projected onto a standard mouse brain template (Allen Mouse Common Coordinate Framework v.3). Edges and nodes are colored according to their anatomical location (Dark Green: Cerebellum, Light Green: Cingulate Area, Red: Hippocampal Formation, Orange: Fiber Tracts, Dark Blue: Olfactory Area, Light Blue: Temporal Association Area, Light Pink: Cerebral nuclei, Purple: Cortical Subplate). Nodes were defined using the DSURQE atlas and restricted to the regions of interest with a significant group difference in brain volumes based on the ENR > STD contrast. Values of bilateral regions were averaged and plotted in both hemispheres. Brain connectivity maps and circular connectogram were generated using NeuroMArVL (http://immersive.erc.monash.edu.au/neuromarvl).

## Discussion

Correlative analyses of MRI data provide evidence of integrated networks of brain regions, both structurally and functionally. These provide non-invasive insights into the organization of the brain at the macroscale, going beyond the analysis of local changes within individual brain regions as revealed by univariate analysis methods (22). Within this framework, ‘structural covariance’, permits the study of inter-individual differences in the organization of brain regional structures within networks that might emerge across a population of individuals in both humans (8, 23) and mice (24, 25). Applying structural covariance and NBS analysis to our individuality model, we observed patterns of structural covariance across the mouse brain that are highly distinct between STD and ENR-housed mice, which may reflect synchronized plastic changes occurring across multiple brain regions over time. Remarkably, within the ENR group itself, the structural covariance matrix for the ‘down’ roamers was similar to the matrix generated for standard-housed mice (Fig. 1 F and G). The structural covariance matrix for those mice with a ‘flat’ RE suggests a much higher degree of inter-regional correlation in comparison to ‘down’ or STD mice, findings confirmed and extended by the NBS analysis. Variation in mouse brain structural covariance in the ENR group suggests that individual behavioral trajectories (as here first approximated by RE, but in reality encompassing the full scope and complexity of behaviors (26) are associated with a certain degree of individualization of brain structural networks at the macroscale and that this largely explains differences observed between standard and ENR mice *per se*. Our data also suggest that RE behavior is supported by widely distributed brain circuitry, in particular the hippocampal formation, cingulate and temporal cortices, white matter tracts and olfactory areas. Consistent with this view, neuroanatomical and/or structural covariance changes within these mouse brain circuits have been previously shown to be associated with spatial memory (27), navigation and cognitive strategies (28) social behaviors (24) and susceptibility or resilience to stress exposure (29) all domains of mouse behavior that may contribute to the individual estimate of RE.

The mice in our study are inbred, thus the genetic background and the “nominal” environment are identical for all mice in the ENR group. Hence, phenotypic variation at the level of brain and behavior must arise from the non-shared component of the environmental factors (‘non-shared environment’). Collectively, our data support a view that a continuum exists between RE and experience-dependent plasticity in brain structure (as measured by *ex vivo* structural MRI). We speculate that exposure to ENR may build on initial variance in mouse brain volumes observable from early in postnatal life (27) and amplifies these differences by providing the opportunity for the development of individual behavior (11, 12, 16, 17). Longitudinal *in vivo* multimodal MRI studies are however required to definitively confirm this view and define the relative contributions of pre-existing individual differences in both functional and structural mouse brain networks as compared to the experience of the shared enriched environment and link these changes to their cellular and molecular correlates.

Because ‘flat’ roamers show stronger structural brain connectivity (based on structural covariance in MRI), it is tempting to speculate that keeping a stable level of territorial coverage has a positive effect on brain networks. In fact, our RE measure has already been used in human studies, providing evidence that greater levels of RE (assessed with a smartphone app) are associated with a greater positive affect and greater hippocampal-striatal functional connectivity (30). Our animal studies allow the generalization of such observations and ultimately will enable identification of the underlying mechanisms at the cellular and molecular levels. Even now, however, our study demonstrates how whole mouse brain structures are distinctively affected by the non-shared component of environmental enrichment and can be linked to stable behavioral trajectories.

## Materials and Methods

### Animals and housing conditions

Female C57BL/6JRj mice, purchased at four weeks of age from Janvier Labs, were randomly divided into standard (STD) (n=10) and enriched (ENR) (n=38) housing conditions. ENR mice were subcutaneously injected into their neck with a glass coated micro transponder (SID 102/A/2; Euro I.D.) under brief isoflurane anesthesia. The shared enriched environment took place in a cage system custom-built to our specifications (PhenoSys GmbH, now marketed as “PhenoSys ColonyRack Individuality 3.0”), consisting of 70 polycarbonate cages (1264C Type II, Tecniplast) that are connected via transparent tunnels and distributed on seven levels. With this cage system, longitudinal tracking of mice is obtained by radio-frequency identification (RFID) antennae, located on connecting tunnels. For this specific experiment, ENR mice only had access to 35 polycarbonate cages in four levels (total area of 1.37 m^2^), equipped with toys and hideouts, that were replaced and rearranged once a week. STD mice were housed in 2 polycarbonate cages (36.5×20.7×14 cm; Type III, Tecniplast) in groups of five animals per cage. All mice were maintained on a 12 hr light/12 hr dark cycle with 55 ± 10% humidity at the animal facility of the Center for Regenerative Therapies Dresden. Food (#V1534; Sniff) and water were provided *ad libitum*.

The experiment was conducted in accordance with the applicable European and national regulations and approved by the local authority (Landesdirektion Sachsen, file number 7/2016 TVT DD24 5131-365-8-SAC).

### Analysis of the tracking data

Antenna contacts, resulting from mice activity, were recorded with the software PhenoSoft Control (PhenoSys GmbH), which identifies the specific antenna and mouse, as well as the time stamp of the antenna contact. As previously described (17), Shannon entropy of the roaming distribution was calculated as 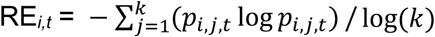, where *i* is the mouse, *j* is the antenna, *k* is the total number of antennae, and *t* is the day. Roaming entropy (RE) quantifies differences in territorial coverage, and because mice are nocturnal animals, only the dark phase of the cycle was used for this analysis. From the nightly mean roaming entropy values, four time-blocks (T1, T2, T3, and T4), of 21 calendar days each, was generated.

### Mixed linear models and repeatability estimation

To investigate whether the observed phenotypic variances are resulting from inter- and/or within-individual variability, we employed generalized linear mixed models in a Bayesian framework, as previously described (12, 31). Briefly, repeatability (R) is the fraction of total variance that can be attributed to interindividual differences, rather than within individual differences, calculated as 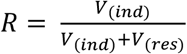, where V_(ind)_ represents the interindividual variances and V_(res)_ the residual, or within individual, variances (32). The behavioral phenotypes from RE were mean-centered and scaled to unity variance. We used time blocks as fixed effect and an interaction between time block and individual identifier as random effect. With a random effect, it was possible to estimate interindividual variances for each time block, while the residual variances were estimated separately. To fit the model, we applied Markov chain Monte Carlo estimation with Gibbs sampling (MCMCglmm R package). A Gaussian error distribution was assumed with weakly informative default priors for the fixed effect (time blocks). An inverse Wishart distribution prior was selected for residual variances. From the posterior (co)variance distributions, we derived the estimates of repeatability and interindividual correlations, using a mode of the posterior density.

### Within enrichment subgroups based on slope patterns

The optimal number of clusters was determined by a silhouette analysis using an unsupervised partitioning method of clustering (k-means), which measures the quality of clustering for each data point(33). All enriched mice were clustered based on mean RE values for each time period (T1, T2, T3, and T4).

A visual assessment of individual behavioral trajectories, based on RE, indicates two distinct patterns: fixed amount of territorial coverage (‘flat’) or behavioral habituation over time (‘down’). After detecting an optimal number of two clusters within our enriched group, we applied slope values, calculated as 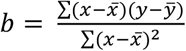, where x are time points (1, 2, 3, and 4, representing time-blocks), y are RE values for each time point, and 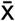 and 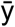 are sample means of the known x’s and the known y’s. The slope values fall into a small distance range: -0.049 up to 0.016. We decided to select up to 15 animals for each subgroup, due to the uncertain behavioral pattern that some mice showed. Finally, 15 animals from the lower spectrum of slope values (−0.049 to -0.008) were grouped as ‘down’ roamers, and 15 animals with slope values ranging from -0.003 up to 0.004 were classified as ‘flat’ roamers. Excluded from the subgroup analysis are 2 mice with in-between slope values and 6 mice with higher than 0.006 slope values.

### Tissue preparation for ex vivo magnetic resonance imaging (MRI)

All animals were culled at 17 weeks of age by cardiac perfusion performed on site at the Kempermann Lab, Center for Regenerative Therapies Dresden, prior to shipping to King’s College London (KCL). The details of the perfusion protocol have been published elsewhere (34, 35). Briefly, mice were anesthetized with sodium pentobarbital (Narcoren 16 g/100ml; 5 μl/g; i.p.) and intracardially perfused with 30 mL of 0.1M phosphate buffered saline (PBS), containing 10 U/mL heparin, followed by 30 mL of 4% (v/v) paraformaldehyde (PFA). Post-perfusion, the mouse was decapitated, and the skin, ears and lower jaw removed, and the brain left within the cranium. This is done to minimize deformations due to dissection. Each brain is then first incubated for 24 hours in 4% PFA solution and then placed in 0.1M PBS containing 0.05% (w/v) sodium azide. Samples were then shipped to KCL and stored at 4°C in this solution for 4 weeks prior to *ex vivo* MR image acquisition to allow tissue rehydration.

### Tissue preparation for immunohistochemistry

Brains in the skulls were shipped back to Dresden from the King’s College London (KCL). At the German Center for Neurodegenerative Diseases (DZNE) Dresden, brains were dissected from the skulls, incubated in 30 % sucrose in phosphate buffer for two days and cut into 40 μm coronal sections using a dry-ice-cooled copper block on a sliding microtome (Leica, SM2000R). Sections were stored at 4 °C in cryoprotectant solution (25 % ethyleneglycol, 25 % glycerol in 0.1 M phosphate buffer, pH 7.4). For detection of Doublecortin (DCX) positive cells, the peroxidase method was applied. Free-floating sections were pre-treated with 0.2 M boric acid (pH 9) at 70 °C for one hour as an antigen retrieval method, washed in 1% phosphate buffer (PBS) and then incubated for 12 hours at 4 °C in PBS with 10 % donkey serum (Jackson Immuno Research Labs) and 0.2 % Triton X-100 (Carl Roth). After the protein blocking step, sections were incubated overnight at 4 °C with the primary antibody (goat anti-DCX, 1:250; Santa Cruz Biotechnology) diluted in PBS containing 3 % donkey serum and 0.2 % Triton X-100. Following washing steps, incubation with biotinylated secondary antibody (1:500, Jackson Immuno Research Labs) occurred for 3 h at room temperature. Sections were then incubated in 0.6 % hydrogen peroxide in PBS for 30 min to inhibit endogenous peroxidase activity. Detection was performed using the Vectastain ABC Elite reagent (9 μg/ml of each component, Vector Laboratories, LINARIS) with diaminobenzidine (0.075 mg/ml; Sigma) and 0.04 % nickel chloride as a chromogen. Stained sections were mounted onto glass slides, cleared with Neo-Clear (Millipore) and cover-slipped using Neo-Mount (Millipore). Doublecortin positive cells were counted on every sixth section along the entire rostro-caudal axis of the dentate gyrus using a brightfield microscope (Leica DM 750).

### MR image acquisition

A 9.4T Bruker BioSpec 94/20 horizontal small bore magnet (Bruker Ltd, UK) and a quadrature volume radiofrequency coil (39 mm internal diameter, Rapid Biomedical GmbH, GER) were used for all *ex vivo* MRI acquisitions. Fixed brain samples were placed securely, four at a time, in a custom-made MR-compatible holder and immersed in proton-free susceptibility matching fluid (Fomblin; Solvay, UK). Samples were scanned in a random order, with the operator blinded to treatment group by numerical coding of samples. Scanning was interspersed with phantoms to ensure consistent operation of the scanner. T2-weighted images were acquired using a 3D fast spin-echo (FSE) sequence with the following parameters: effective echo time (TE) 30 ms, repetition time (TR) 3000 ms, field of view (FOV) 25×25×20 mm and acquisition matrix 250×250×200 yielding isotropic voxels of 100 μm^3^, scan time = 5 hours and 44 minutes.

### MR image processing

MR images were visually inspected in native space for artefacts, with no images excluded on this basis. Raw MR Images were converted from the manufacturer’s proprietary format to the NIFTI format and processed using a combination of FSL(36), ANTs(37) and the Quantitative Imaging Tools (QUIT) package written in C++ software utilizing the ITK library, available from *https://github.com/spinicist/QUIT*. The following steps were performed on the T2-FSE anatomical MR images in their native space. A Tukey filter was applied to the FSE MR images in k-space to remove high frequency noise followed by correction for intensity inhomogeneity using the N4 algorithm (38). A study-specific template was then constructed from MR images of n=24 mice randomly selected from the entire dataset, using the *antsMultivariateTemplateConstruction2*.*sh* script with cross-correlation metric and SyN transform (39). The template and individual brains were skull-stripped using the RATS algorithm implemented in QUIT using the *qimask* script(35).

### MR image analyses

#### Atlas based segmentation of regional brain volumes

Group level differences in volume were assessed using a combination of ABS and voxel-wise DBM as in our prior work(25). To enable atlas based segmentation (ABS) analysis of regional brain volumes, the study-specific template was then registered to the Dorr-Steadman-Ullmann-Richards-Qiu-Egan (DSURQE) mouse brain atlas (40 micron) (https://wiki.mouseimaging.ca/display/MICePub/Mouse+Brain+Atlases). This atlas has 182 individual mouse brain structures defined (including left and right labels for most structures, giving 356 labels in total). The T2-weighted 3D FSE images for all study subjects were then non-linearly registered to the study template using the *antsRegistration-SyN*.*sh* script. For the DSURQE atlas, the inverse transforms from the atlas to the study template and from the study template to each subject, were applied to calculate the brain and atlas based ROI volumes for each subject (40, 41). After careful checking of atlas label alignment to each individual subject’s MR images, we automatically extracted volumes for the 182 ROIs comprising the DSURQE atlas, merging the left and right labels for each ROI. Total brain volume was calculated from the summation of the individual atlas ROI volumes. Both absolute volumes (mm^3^) and relative volumes were compared to assess brain scaling. Relative volumes were calculated as a percentage of total brain volume i.e. ([(brain region volume)/(whole brain volume) × 100]. Whilst others and we have used this approach there are caveats. Specifically, whilst some brain regions scale linearly to total brain volume, such as the hippocampus, other regions, such as the cerebellum, may not (42). Hence it is important to compare both absolute and relative volumes in assessing group differences. Group-level differences in either absolute or relative volumes, were compared between STD and ENR mice using multiple t-tests (2-tailed, unequal variance assumed) with α=0.05 using R-project (v4.0). The resulting p-values were subsequently corrected for multiple comparisons using the false discovery rate (FDR) at 5% (q<0.05) using Prism software (v8.4.2; GraphPad, La Jolla, CA, USA). To calculate the magnitude and direction of volume change for each region between groups, effect sizes for each brain region were calculated using the standardized mean difference (effect size = (μ _[ENR]_ -μ _[STD]_ / σ _STD_); measured in units of standard deviation.

#### Deformation based morphometry

Jacobian determinant maps were calculated from the inverse warp fields in standard space using the *CreateJacobianDeterminantImage* script (ANTs) and log-scaled to allow voxel-wise estimation of apparent volume change via deformation based morphometry (DBM) (*25*). To compare local volume differences between STD and ENR mice, voxel-based nonparametric statistics were performed on the log-transformed Jacobian determinant maps using FSL randomise as previously described(25). The resulting F-statistic maps were corrected for multiple comparisons using the family-wise error rate (FWE, p < 0.05). Data in the manuscript are also shown at p<0.05 uncorrected for multiple comparisons. DBM analyses were run with and without total brain volume as a covariate of no interest, to check for potential global scaling effects.

#### Correlation between RE and local volume changes

The slope of the RE metric, calculated across all time blocks was regressed against log-Jacobians of each individual at each voxel in the ENR cohort only. The resultant correlation map was then thresholded using the family-wise error rate (FWE, p < 0.05). Data in the manuscript are also shown at p<0.05 uncorrected for multiple comparisons.

#### Mouse brain structural covariance

Following the atlas-based approach, we explored patterns of structural covariance using relative brain volumes from 182 predefined structures in the DSURQE atlas (10). We computed the Pearson correlation coefficient between all relative structure volumes, resulting in a 182×182 matrix of correlations representing the structural covariance network, for both standard housing and enriched housing groups. We repeated the same analysis considering the within-enrichment subgroups ‘flat’ and ‘down’ roamers. The 182 structures are ordered following hierarchical clustering of the enriched-housed correlation matrix, with the same order applied for all matrices.

#### Network based statistic (NBS) analysis

Network based statistics (NBS) is a powerful statistical method for identifying a statistically significant cluster of connections indicating differences between groups on intermediate network scales (43). As described in previous functional studies (44, 45), NBS are computed using the following steps: (i) identify all connections (pairs of nodes) that are different between groups beyond a particular t-value (called primary threshold), (ii) select the largest contiguous cluster of these connections, and (iii) validate the cluster’s significance by permutation testing. In permutation testing an empirical null distribution of the largest cluster size is generated by conducting the first two NBS steps on resampled group membership data 10,000 times. The returned subnetwork is statistically significant at a family wise error (FWE) corrected value of p<0.05.

Although the network needs to be considered as a whole, the extent of the returned network can be varied using a different primary threshold. This adjusts the extremity of deviation in a connection between groups required, before it is considered for inclusion in the NBS result. NBS returns a single p-value, which represents the likelihood that the sub-network (also called component) is due to a true effect in the data. This approach measures the entire cluster of returned connections but does not identify the contribution of each connection independently. The NBS procedure was carried out for the correlation matrices with a primary threshold of t=3.1 (which corresponds to p=0.001).

As part of an additional analysis to see if cortical regions that are susceptible to thickness reductions are more interconnected the NBS procedure was carried out for STD and ENR on a subgroup of ROIs consisting of all significantly changed regions between STD and ENR.

## References

1. R. Plomin, D. Daniels, Why are children in the same family so different from one another? Behav Brain Sci 10, 1–16 (1987).

2. F. Morgante, P. Sørensen, D. A. Sorensen, C. Maltecca, T. F. C. Mackay, Genetic Architecture of Micro-Environmental Plasticity in Drosophila melanogaster. Sci Rep 5, 9785 (2015).

3. B. Draganski, F. Kherif, A. Lutti, Computational anatomy for studying use-dependant brain plasticity. Front. Hum. Neurosci. 8 (2014).

4. M. Taubert, et al., Dynamic properties of human brain structure: learning-related changes in cortical areas and associated fiber connections. J Neurosci 30, 11670–11677 (2010).

5. A. D. Nostro, V. I. Müller, A. T. Reid, S. B. Eickhoff, Correlations Between Personality and Brain Structure: A Crucial Role of Gender. Cereb Cortex 27, 3698–3712 (2017).

6. R. Kanai, T. Feilden, C. Firth, G. Rees, Political orientations are correlated with brain structure in young adults. Curr Biol 21, 677–680 (2011).

7. F. Melozzi, et al., Individual structural features constrain the mouse functional connectome. Proc Natl Acad Sci USA 116, 26961–26969 (2019).

8. A. Alexander-Bloch, J. N. Giedd, E. Bullmore, Imaging structural co-variance between human brain regions. Nat Rev Neurosci 14, 322–336 (2013).

9. M. Pagani, A. Bifone, A. Gozzi, Structural covariance networks in the mouse brain. Neuroimage 129, 55–63 (2016).

10. Y. Yee, et al., Structural covariance of brain region volumes is associated with both structural connectivity and transcriptomic similarity. Neuroimage 179, 357–372 (2018).

11. J. C. Körholz, et al., Selective increases in inter-individual variability in response to environmental enrichment in female mice. eLife 7, e35690 (2018).

12. S. Zocher, et al., Early-life environmental enrichment generates persistent individualized behavior in mice. Sci. Adv. 6, eabb1478 (2020).

13. T.-Y. Zhang, et al., Environmental enrichment increases transcriptional and epigenetic differentiation between mouse dorsal and ventral dentate gyrus. Nat Commun 9, 298 (2018).

14. J. Scholz, R. Allemang-Grand, J. Dazai, J. P. Lerch, Environmental enrichment is associated with rapid volumetric brain changes in adult mice. NeuroImage 109, 190–198 (2015).

15. Y. Stern, et al., Whitepaper: Defining and investigating cognitive reserve, brain reserve, and brain maintenance. Alzheimer’s &amp; Dementia 16, 1305–1311 (2020).

16. G. Kempermann, Environmental enrichment, new neurons and the neurobiology of individuality. Nat Rev Neurosci 20, 235–245 (2019).

17. J. Freund, et al., Emergence of Individuality in Genetically Identical Mice. Science 340, 756–759 (2013).

18. M. C. Diamond, et al., Increases in cortical depth and glia numbers in rats subjected to enriched environment. J. Comp. Neurol. 128, 117–125 (1966).

19. G. Gong, Y. He, Z. J. Chen, A. C. Evans, Convergence and divergence of thickness correlations with diffusion connections across the human cerebral cortex. Neuroimage 59, 1239–1248 (2012).

20. J. M. Segall, et al., Correspondence between structure and function in the human brain at rest. Front Neuroinform 6, 10 (2012).

21. J. H. Steiger, Tests for comparing elements of a correlation matrix. Psychological Bulletin 87, 245–251 (1980).

22. E. Bullmore, O. Sporns, Complex brain networks: graph theoretical analysis of structural and functional systems. Nat Rev Neurosci 10, 186–198 (2009).

23. A. C. Evans, Networks of anatomical covariance. Neuroimage 80, 489–504 (2013).

24. M. R. Bruce, et al., Sexually dimorphic neuroanatomical differences relate to ASD-relevant behavioral outcomes in a maternal autoantibody mouse model. Mol Psychiatry (2021) https://doi.org/10.1038/s41380-021-01215-w.

25. F. S. Mueller, et al., Behavioral, neuroanatomical, and molecular correlates of resilience and susceptibility to maternal immune activation. Mol Psychiatry 26, 396–410 (2021).

26. J. Freund, et al., Association between exploratory activity and social individuality in genetically identical mice living in the same enriched environment. Neuroscience 309, 140–152 (2015).

27. J. P. Lerch, et al., Maze training in mice induces MRI-detectable brain shape changes specific to the type of learning. Neuroimage 54, 2086–2095 (2011).

28. A. Qiu, et al., Morphology and microstructure of subcortical structures at birth: A large-scale Asian neonatal neuroimaging study. NeuroImage 65, 315–323 (2013).

29. C. Anacker, et al., Neuroanatomic Differences Associated With Stress Susceptibility and Resilience. Biol Psychiatry 79, 840–849 (2016).

30. A. S. Heller, et al., Association between real-world experiential diversity and positive affect relates to hippocampal-striatal functional connectivity. Nat Neurosci 23, 800–804 (2020).

31. Y. Fong, H. Rue, J. Wakefield, Bayesian inference for generalized linear mixed models. Biostatistics 11, 397–412 (2010).

32. N. J. Dingemanse, N. A. Dochtermann, Quantifying individual variation in behaviour: mixed-effect modelling approaches. J Anim Ecol 82, 39–54 (2013).

33. L. Kaufman, P. J. Rousseeuw, Finding groups in data: an introduction to cluster analysis (Wiley, 2005).

34. J. Richetto, et al., Genome-Wide Transcriptional Profiling and Structural Magnetic Resonance Imaging in the Maternal Immune Activation Model of Neurodevelopmental Disorders. Cereb Cortex 27, 3397–3413 (2017).

35. T. C. Wood, et al., Whole-brain ex-vivo quantitative MRI of the cuprizone mouse model. PeerJ 4, e2632 (2016).

36. M. Jenkinson, C. F. Beckmann, T. E. J. Behrens, M. W. Woolrich, S. M. Smith, FSL. Neuroimage 62, 782–790 (2012).

37. B. B. Avants, et al., A reproducible evaluation of ANTs similarity metric performance in brain image registration. Neuroimage 54, 2033–2044 (2011).

38. N. J. Tustison, et al., N4ITK: improved N3 bias correction. IEEE Trans Med Imaging 29, 1310–1320 (2010).

39. B. B. Avants, et al., The optimal template effect in hippocampus studies of diseased populations. NeuroImage 49, 2457–2466 (2010).

40. N. Doostdar, et al., Global brain volume reductions in a sub-chronic phencyclidine animal model for schizophrenia and their relationship to recognition memory. J Psychopharmacol 33, 1274–1287 (2019).

41. W.-L. Kuan, et al., Systemic α-synuclein injection triggers selective neuronal pathology as seen in patients with Parkinson’s disease. Mol Psychiatry 26, 556–567 (2021).

42. C. Mankiw, et al., Allometric Analysis Detects Brain Size-Independent Effects of Sex and Sex Chromosome Complement on Human Cerebellar Organization. J. Neurosci. 37, 5221–5231 (2017).

43. A. Zalesky, A. Fornito, E. T. Bullmore, Network-based statistic: identifying differences in brain networks. Neuroimage 53, 1197–1207 (2010).

44. D. Geisler, et al., Altered global brain network topology as a trait marker in patients with anorexia nervosa. Psychol. Med. 50, 107–115 (2020).

45. S. Ehrlich, et al., Reduced functional connectivity in the thalamo-insular subnetwork in patients with acute anorexia nervosa: Functional Connectivity in Anorexia Nervosa. Hum. Brain Mapp. 36, 1772–1781 (2015).

