## Supplemental Material for "Individual behavioral trajectories shape whole-brain connectivity in mice"

A visual assessment of individual behavioural trajectories, based on RE, indicates two distinct patterns: fixed amount of territorial coverage (‘flat’) or behavioural habituation over time (‘down’). After detecting an optimal number of two clusters within our enriched group, we applied slope values, calculated as $b = \frac{\Sigma(x-\bar{x})(y-\bar{y})}{\Sigma(x-\bar{x})^2}$, where x are time points (1, 2, 3, and 4, representing time-blocks), y are RE values for each time point, and $\bar{x}$ and $\bar{y}$ are sample means of the known x's and the known y's. The slope values fall into a small distance range: -0.049 up to 0.016. We decided to select up to 15 animals for each subgroup, due to the uncertain behavioural pattern that some mice showed. Finally, 15 animals from the lower spectrum of slope values (-0.049 to -0.008) were grouped as ‘down’ roamers, and 15 animals with slope values ranging from -0.003 up to 0.004 were classified as ‘flat’ roamers. Excluded from the subgroup analysis are 2 mice with in-between slope values and 6 mice with higher than 0.006 slope values.

### Supplementary Figures

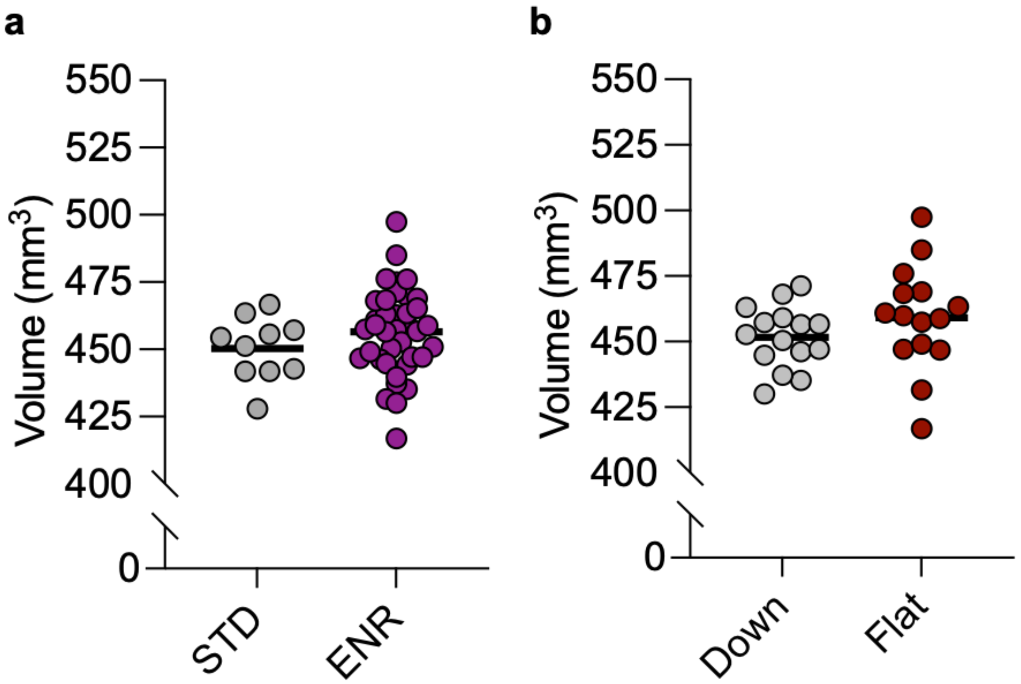


**Figure S1:** Total brain volumes (as measured by ex vivo structural MRI) do not differ significantly when comparing either (a) STD vs. ENR-housed mice or (b) Down vs. Flat sub-groups of ENR-housed mice. Each data point indicates the whole brain volume (in $mm^3$) for an individual mouse in each group. Solid horizontal line indicates the group mean.

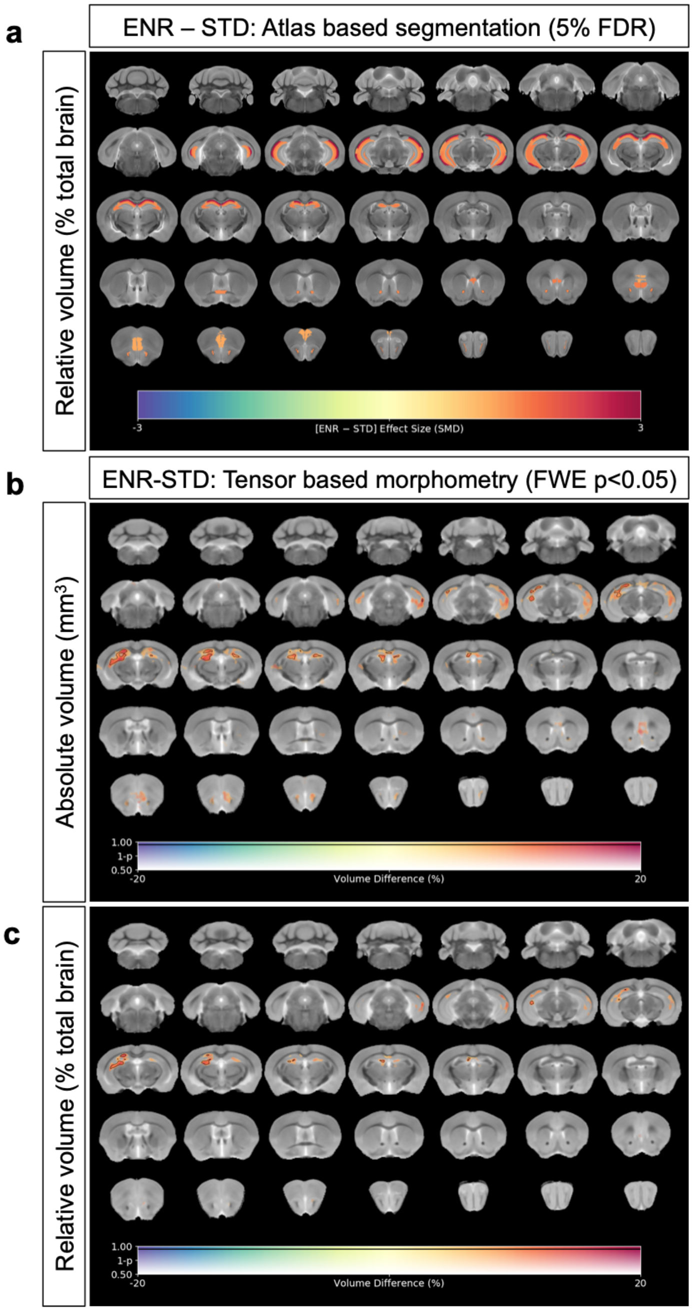


**Figure S2:** (a) Map of regional differences in relative volume (% total brain volume) extracted using the DSURQE mouse brain atlas comparing ENR mice to those in standard (STD) housing. Only atlas regions of interest (ROI) that survive multiple comparisons correction are shown (False Discovery Rate, 5%). Data shown for each statistically significant ROI is the standardized mean difference (SMD) comparing ENR – STD housed mice. Warm colors indicate regional volume increases, whilst cold colors indicate regional volume decreases in ENR-mice as compared to those in STD housing. (b-c) Complementary voxel-wise tensor based morphometry (TBM) to map localized neuroanatomical differences between ENR-STD mice. Data shown in (b, c) are the percentage difference in log-scaled Jacobian determinants either without correction for total brain volume (b; absolute volumes) or with correction for total brain volume (c; relative volumes) between ENR and STD-housed mice. Voxels with solid black contours survive a stringent multiple comparisons correction (Family Wise Error $p<0.05$) all other voxels reflect an exploratory threshold of $p<0.05$ uncorrected for multiple comparisons. Warm colors indicate voxels with increased volume comparing ENR to STD mice, whilst cold colors indicate voxels with reduced volume.

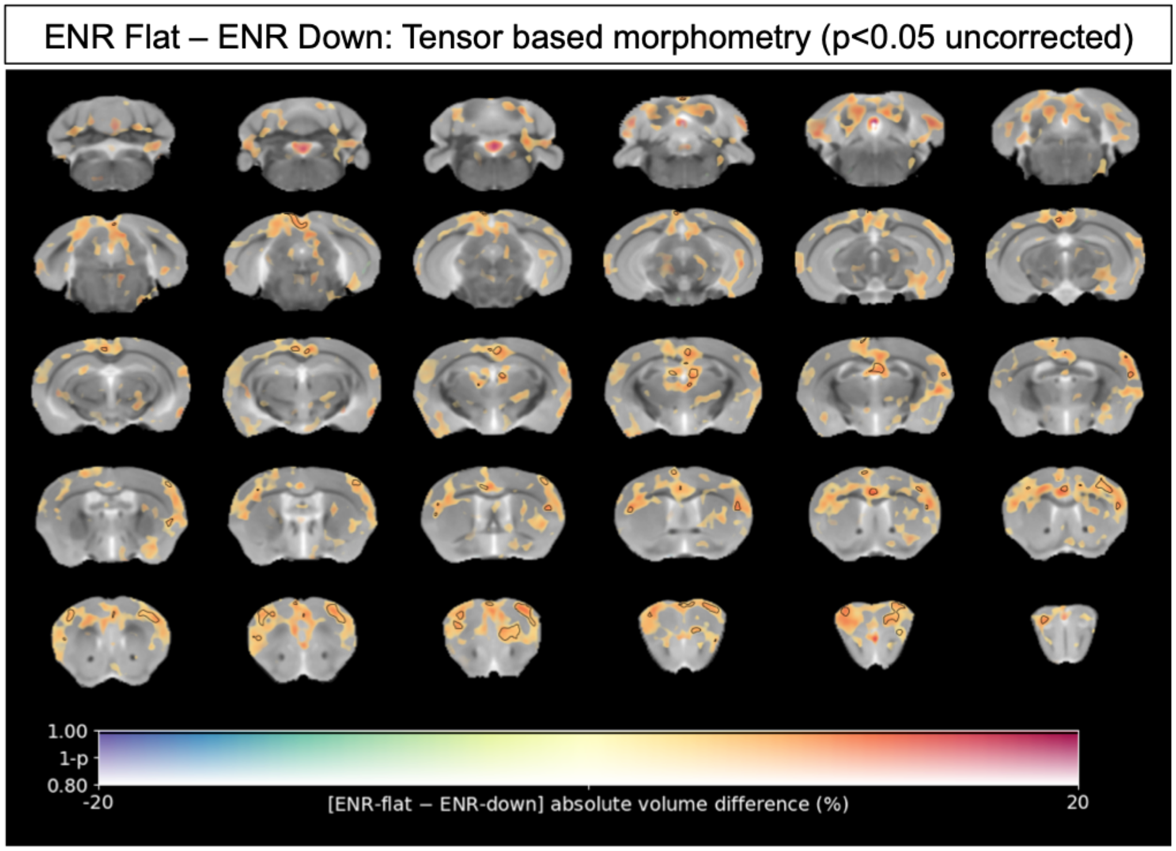


**Figure S3**: Voxel-wise tensor based morphometry (TBM) to map localized neuroanatomical differences between Flat and Down subgroups of ENR-housed mice. Data shown are the percentage difference in log-scaled Jacobian determinants without correction for total brain volume (absolute volumes) between Flat and Down subgroups of ENR-housed mice. There were no statistically significant localized volume differences between the groups after a stringent correction for multiple comparisons (Family Wise Error $p<0.05$). Voxels with solid black contours therefore represent apparent local volume differences between Flat and Down mice at an exploratory threshold of $p<0.05$ uncorrected for multiple comparisons. Warm colors indicate voxels with apparent increases volume comparing Flat to Down mice, whilst cold colors indicate voxels with apparent reductions in volume.

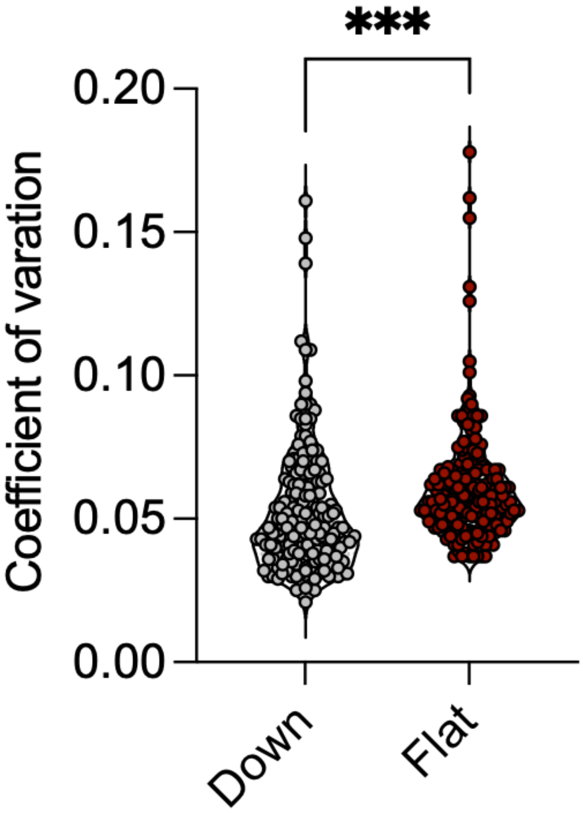


**Figure S4:** The violin plots with overlaid data points show increased overall coefficient variance in the volume of mouse brain regions in Flat compared to Down sub-groups of ENR-housed mice. Individual data points represent the coefficient of variation (CV) for each of 182 regions of interest in the DSURQE mouse brain atlas. ***p<0.001 Mann-Whitney U test comparing the average of the CV across all 182 ROIs in the DSURQE mouse brain atlas, to yield an average variability measure for each group.

### Supplemental Table S1

| Absolute volume (mm3) | STR | | ENR | | | | |
|---|---|---|---|---|---|---|---|
| Atlas region of interest | Mean | SD | Mean | SD | Effect size | p-value | q-value |
| CA10r | 2,01427752 | 0,05518613 | 2,16316227 | 0,08775506 | 2,70 | 1,0621E-06 | 0,0002 |
| CA3Rad | 1,8663607 | 0,0501575 | 1,9770504 | 0,07136835 | 2,21 | 1,6876E-05 | 0,0010 |
| CA1Py | 1,01038984 | 0,03383358 | 1,08495587 | 0,04499654 | 2,20 | 1,7227E-05 | 0,0010 |
| Cingulate cortex: area 25 | 0,55209421 | 0,02338139 | 0,60155064 | 0,04484816 | 2,12 | 5,0712E-05 | 0,0017 |
| subependymale zone / rhinocele | 0,05391439 | 0,0022136 | 0,05849994 | 0,00280695 | 2,07 | 3,6106E-05 | 0,0016 |
| CA2Rad | 0,4964123 | 0,01704977 | 0,53057872 | 0,02519475 | 2,00 | 5,5843E-05 | 0,0017 |
| CA2Py | 0,18176779 | 0,00646279 | 0,1943868 | 0,00921149 | 1,95 | 7,3108E-05 | 0,0019 |
| SLu | 0,62483991 | 0,01573946 | 0,65488075 | 0,02795656 | 1,91 | 0,0001 | 0,0032 |
| anterior commissure: pars anterior | 1,13732077 | 0,0355595 | 1,1997492 | 0,03964941 | 1,76 | 0,0002 | 0,0039 |
| CA1Rad | 2,46686483 | 0,09666775 | 2,63614309 | 0,09761858 | 1,75 | 0,0002 | 0,0039 |
| CA20r | 0,46745264 | 0,01884719 | 0,49925226 | 0,0225488 | 1,69 | 0,0003 | 0,0051 |
| CA3Py Inner | 0,11296331 | 0,0045581 | 0,12032353 | 0,00385686 | 1,61 | 0,0005 | 0,0070 |
| Medial orbital cortex | 1,83526921 | 0,07235229 | 1,94843435 | 0,10961163 | 1,56 | 0,0008 | 0,0104 |
| LMol | 2,35432494 | 0,09208074 | 2,49540701 | 0,09883332 | 1,53 | 0,0007 | 0,0100 |
| CA3Py Outer | 0,9416147 | 0,024131 | 0,97837139 | 0,04615404 | 1,52 | 0,0018 | 0,0209 |
| Dorsal tenia tecta | 1,0259587 | 0,04808944 | 1,09766288 | 0,08184744 | 1,49 | 0,0016 | 0,0192 |
| GrDG | 1,22102802 | 0,05521758 | 1,29447134 | 0,04236333 | 1,33 | 0,0021 | 0,0223 |
| subiculum | 3,43229402 | 0,13268705 | 3,60060252 | 0,11458908 | 1,27 | 0,0029 | 0,0296 |
| olfactory peduncle | 2,54050135 | 0,06655028 | 2,62269457 | 0,09037459 | 1,24 | 0,0047 | 0,0409 |
| PoDG | 0,51477325 | 0,02646059 | 0,54738331 | 0,01828178 | 1,23 | 0,0035 | 0,0333 |
| Anterior olfactory nucleus | 2,2415151 | 0,05999757 | 2,31485342 | 0,07596006 | 1,22 | 0,0047 | 0,0409 |
| MoDG | 4,24515853 | 0,18098194 | 4,45254727 | 0,15775464 | 1,15 | 0,0057 | 0,0476 |
| Olfactory bulb: internal plexiform layer | 0,72914169 | 0,01625352 | 0,75024498 | 0,03376129 | 1,30 | 0,0085 | 0,06 |
| Olfactory bulb: mitral cell layer | 1,01905838 | 0,02380348 | 1,04995752 | 0,04678252 | 1,30 | 0,0072 | 0,06 |
| Olfactory bulb: external plexiform layer | 7,04048148 | 0,18073162 | 7,22920799 | 0,320104 | 1,04 | 0,0217 | 0,14 |
| Olfactory bulb: granule cell layer | 4,46834483 | 0,14750605 | 4,61880334 | 0,20000592 | 1,02 | 0,0160 | 0,11 |
| fimbria | 3,12303105 | 0,12232506 | 3,23899571 | 0,20744904 | 0,95 | 0,0329 | 0,18 |
| bed nucleus of stria terminalis | 1,26061638 | 0,04037353 | 1,29878652 | 0,06287226 | 0,95 | 0,0290 | 0,17 |
| nucleus accumbens | 4,26586038 | 0,17030186 | 4,42679712 | 0,19504048 | 0,95 | 0,0204 | 0,13 |
| Accessory olfactory bulb: granule cell layer | 0,18509417 | 0,01037137 | 0,19489173 | 0,00947742 | 0,94 | 0,0178 | 0,12 |
| lateral septum | 2,97715899 | 0,11963679 | 3,08958046 | 0,16012661 | 0,94 | 0,0245 | 0,15 |
| corpus callosum | 12,7111975 | 0,36218664 | 13,0487911 | 0,61886267 | 0,93 | 0,0361 | 0,19 |

| | | | | | | | |
|---|---|---|---|---|---|---|---|
| Temporal association area | 2,55289942 | 0,08593477 | 2,62895536 | 0,11614069 | 0,89 | 0,0332 | 0,18 |
| Cingulum | 0,8812946 | 0,04054711 | 0,91417438 | 0,04951053 | 0,81 | 0,0444 | 0,23 |
| Cingulate cortex: area 32 | 2,54245002 | 0,17650977 | 2,67580692 | 0,14518729 | 0,76 | 0,0474 | 0,24 |
| lobule 8 white matter | 0,10624584 | 0,0076673 | 0,10038173 | 0,00760907 | -0,76 | 0,0490 | 0,24 |
| trunk of lobules 6-8 white matter | 0,13381498 | 0,007852 | 0,12557509 | 0,00922248 | -1,05 | 0,0117 | 0,08 |
| Medial preoptic nucleus | 0,17233697 | 0,01156502 | 0,17847637 | 0,0122732 | 3,6 | 0,1613 | 0,4940 |
| Cingulate cortex: area 24a | 1,90104315 | 0,0978624 | 1,96585005 | 0,12319167 | 3,4 | 0,0962 | 0,3647 |
| cerebral aqueduct | 0,37384806 | 0,03020671 | 0,3860326 | 0,05731333 | 3,3 | 0,3685 | 0,6952 |
| Cingulate cortex: area 24b | 1,71043528 | 0,10595067 | 1,76450039 | 0,09083885 | 3,2 | 0,1640 | 0,4940 |
| Secondary somatosensory cortex | 5,92902337 | 0,24093743 | 6,11428374 | 0,29981443 | 3,1 | 0,0560 | 0,2531 |
| thalamus | 17,4494982 | 0,91513898 | 17,9584009 | 1,00996474 | 2,9 | 0,1463 | 0,4843 |
| posterior commissure | 0,15226026 | 0,01340764 | 0,15666735 | 0,01398752 | 2,9 | 0,3743 | 0,6952 |
| Ectorhinal cortex | 2,63406186 | 0,10297924 | 2,70862799 | 0,13334948 | 2,8 | 0,0727 | 0,3070 |
| CA30r | 3,00646428 | 0,09861185 | 3,08917209 | 0,16118712 | 2,8 | 0,0537 | 0,2531 |
| Frontal cortex: area 3 | 0,69976138 | 0,03032205 | 0,71862448 | 0,03121887 | 2,7 | 0,1032 | 0,3730 |
| Cingulate cortex: area 29c | 1,9878155 | 0,09307054 | 2,04033732 | 0,10769323 | 2,6 | 0,1445 | 0,4843 |
| Frontal association cortex | 7,2191934 | 0,28728142 | 7,40339879 | 0,3366061 | 2,6 | 0,1012 | 0,3730 |
| Dorsolateral orbital cortex | 0,92635734 | 0,0375911 | 0,9499246 | 0,04624122 | 2,5 | 0,1120 | 0,3919 |
| paraflocculus white matter | 0,18092579 | 0,01686457 | 0,18550801 | 0,01587575 | 2,5 | 0,4524 | 0,7447 |
| Ventral orbital cortex | 1,58245349 | 0,04948909 | 1,62217603 | 0,07398733 | 2,5 | 0,0570 | 0,2531 |
| internal capsule | 3,18629639 | 0,09970736 | 3,2647789 | 0,17756122 | 2,5 | 0,0776 | 0,3070 |
| Accessory olfactory bulb: glomerular, external plexiform and mitral cell layer | 0,37123059 | 0,02591541 | 0,38021151 | 0,02612071 | 2,4 | 0,3466 | 0,6932 |
| fourth ventricle | 0,72899465 | 0,05693411 | 0,74603472 | 0,06174412 | 2,3 | 0,4211 | 0,7413 |
| striatum | 20,0131082 | 0,591263 | 20,4749143 | 0,7855687 | 2,3 | 0,0559 | 0,2531 |
| mammilothalamic tract | 0,24842823 | 0,00988463 | 0,25407037 | 0,01465572 | 2,3 | 0,1657 | 0,4940 |
| fundus of striatum | 0,17285644 | 0,00570742 | 0,1767277 | 0,00861176 | 2,2 | 0,1045 | 0,3730 |
| globus pallidus | 3,31253517 | 0,17493441 | 3,38644277 | 0,19658754 | 2,2 | 0,2645 | 0,6173 |
| Lateral parietal association cortex | 0,20797595 | 0,01139331 | 0,21259939 | 0,01190519 | 2,2 | 0,2763 | 0,6361 |
| cerebellar peduncle: superior | 1,0377713 | 0,05697723 | 1,06069938 | 0,06731862 | 2,2 | 0,2923 | 0,6361 |
| periaqueductal grey | 3,83602553 | 0,25775779 | 3,92038122 | 0,25800584 | 2,2 | 0,3727 | 0,6952 |
| Secondary auditory cortex: ventral area | 1,47096045 | 0,05935683 | 1,50319571 | 0,07391186 | 2,2 | 0,1659 | 0,4940 |
| pre-para subiculum | 2,17634662 | 0,11924844 | 2,22291403 | 0,1186198 | 2,1 | 0,2898 | 0,6361 |
| lobules 4-5 white matter | 0,50429782 | 0,05040839 | 0,51492879 | 0,03816364 | 2,1 | 0,5459 | 0,7866 |
| Cingulate cortex: area 24a' | 0,85999304 | 0,05440475 | 0,87810997 | 0,04838012 | 2,1 | 0,3555 | 0,6952 |
| Olfactory bulb: glomerular layer | 4,66332682 | 0,12351165 | 4,76031891 | 0,21794782 | 2,1 | 0,0772 | 0,3070 |
| Primary somatosensory cortex | 4,08076772 | 0,15209871 | 4,16461115 | 0,19695192 | 2,1 | 0,1638 | 0,4940 |
| Primary somatosensory cortex: upper lip region | 5,18726894 | 0,20698934 | 5,29374175 | 0,27422327 | 2,1 | 0,1950 | 0,5311 |
| Perirhinal cortex | 2,41352456 | 0,10352341 | 2,46292645 | 0,12939793 | 2,0 | 0,2209 | 0,5583 |
| midbrain | 14,1579817 | 0,73111123 | 14,4271006 | 0,90182598 | 1,9 | 0,3391 | 0,6882 |

| | | | | | | | |
|---|---|---|---|---|---|---|---|
| flocculus white matter | 0,0487534 | 0,00515906 | 0,04967679 | 0,00368693 | 1,9 | 0,6052 | 0,8278 |
| stria medullaris | 0,73850029 | 0,03582067 | 0,75240964 | 0,0396556 | 1,9 | 0,3021 | 0,6410 |
| fornix | 0,73970743 | 0,02740058 | 0,75354603 | 0,03440379 | 1,9 | 0,1967 | 0,5311 |
| colliculus: superior | 8,89178222 | 0,54828619 | 9,05114422 | 0,56205208 | 1,8 | 0,4292 | 0,7413 |
| medial septum | 1,40736152 | 0,04627341 | 1,43215367 | 0,05584848 | 1,8 | 0,1683 | 0,4940 |
| Secondary motor cortex | 6,15074859 | 0,22022175 | 6,25852434 | 0,25918782 | 1,8 | 0,2036 | 0,5369 |
| lobule 10: nodulus | 1,20613629 | 0,05936701 | 1,22632538 | 0,08560801 | 1,7 | 0,3975 | 0,7234 |
| interpeduncular nucleus | 0,26756328 | 0,02078581 | 0,27202743 | 0,02142014 | 1,7 | 0,5575 | 0,7866 |
| Medial parietal association cortex | 0,45298292 | 0,02234849 | 0,46048538 | 0,02457583 | 1,7 | 0,3696 | 0,6952 |
| Caudomedial entorhinal cortex | 4,94266667 | 0,29555427 | 5,02260489 | 0,22450026 | 1,6 | 0,4411 | 0,7413 |
| cerebral peduncle | 2,44676273 | 0,10122792 | 2,485946 | 0,11127536 | 1,6 | 0,3029 | 0,6410 |
| Medial entorhinal cortex | 0,67468408 | 0,04378078 | 0,68473928 | 0,02475037 | 1,5 | 0,5005 | 0,7720 |
| lateral olfactory tract | 1,54079049 | 0,04274479 | 1,56364141 | 0,06581162 | 1,5 | 0,1984 | 0,5311 |
| basal forebrain | 5,59848396 | 0,14334352 | 5,68121403 | 0,23605656 | 1,5 | 0,1763 | 0,5013 |
| Insular region: not subdivided | 6,59012761 | 0,11967816 | 6,68259481 | 0,31296002 | 1,4 | 0,1521 | 0,4940 |
| hypothalamus | 10,9106573 | 0,35704219 | 11,0634824 | 0,51458915 | 1,4 | 0,2893 | 0,6361 |
| Primary somatosensory cortex: jaw region | 0,65840991 | 0,03104769 | 0,66756932 | 0,0315505 | 1,4 | 0,4217 | 0,7413 |
| nucleus interpositus | 0,43600197 | 0,02088932 | 0,44188298 | 0,02490033 | 1,3 | 0,4583 | 0,7447 |
| Cingulate cortex: area 24b' | 0,59455648 | 0,02918769 | 0,60224491 | 0,03277511 | 1,3 | 0,4811 | 0,7618 |
| Claustrum: dorsal part | 0,21846284 | 0,00774913 | 0,22126059 | 0,01007479 | 1,3 | 0,3548 | 0,6952 |
| Secondary visual cortex: lateral area | 2,45818256 | 0,09976974 | 2,4895152 | 0,10027101 | 1,3 | 0,3921 | 0,7209 |
| colliculus: inferior | 5,82208212 | 0,3262345 | 5,89512378 | 0,34438359 | 1,3 | 0,5431 | 0,7866 |
| third ventricle | 1,16627278 | 0,04327209 | 1,18058004 | 0,07581163 | 1,2 | 0,4440 | 0,7413 |
| lobules 4-5: culmen (ventral and dorsal) | 4,32927925 | 0,24045616 | 4,3820106 | 0,27119836 | 1,2 | 0,5570 | 0,7866 |
| Primary motor cortex | 7,18949476 | 0,34293389 | 7,2770013 | 0,34747835 | 1,2 | 0,4856 | 0,7618 |
| Medial amygdala | 1,32441574 | 0,03799423 | 1,34030755 | 0,06848915 | 1,2 | 0,3403 | 0,6882 |
| Cingulate cortex: area 29a | 0,71487043 | 0,03791494 | 0,72339391 | 0,04172463 | 1,2 | 0,5450 | 0,7866 |
| stria terminalis | 0,9321271 | 0,02199914 | 0,94288824 | 0,04487982 | 1,2 | 0,2936 | 0,6361 |
| Secondary visual cortex: mediolateral area | 1,01070724 | 0,04869517 | 1,02192534 | 0,05683259 | 1,1 | 0,5407 | 0,7866 |
| Secondary auditory cortex: dorsal area | 1,30298054 | 0,0555765 | 1,31737634 | 0,0536628 | 1,1 | 0,4753 | 0,7618 |
| ventral tegmental decussation | 0,12947959 | 0,0105305 | 0,1309032 | 0,01121997 | 1,1 | 0,7129 | 0,8709 |
| paraflocculus (PFL) | 2,17826826 | 0,31264972 | 2,201773 | 0,34206896 | 1,1 | 0,8385 | 0,9382 |
| Amygdalopiriform transition area | 1,02235504 | 0,06753632 | 1,0326578 | 0,05755216 | 1,0 | 0,6659 | 0,8417 |
| Ventral intermediate entorhinal cortex | 1,01944783 | 0,07340129 | 1,02943246 | 0,04888487 | 1,0 | 0,6916 | 0,8611 |
| pons | 17,7819362 | 0,66831866 | 17,9548469 | 0,92533239 | 1,0 | 0,5127 | 0,7783 |
| Primary auditory cortex | 1,34553881 | 0,04939325 | 1,35859015 | 0,05737484 | 1,0 | 0,4832 | 0,7618 |
| Dorsal nucleus of the endopiriform | 1,50786665 | 0,03023466 | 1,52210967 | 0,0561427 | 0,9 | 0,2902 | 0,6361 |
| Posteromedial cortical amygdaloid area | 1,32267343 | 0,07444003 | 1,33498741 | 0,07570883 | 0,9 | 0,6498 | 0,8341 |

| | | | | | | | |
|---|---|---|---|---|---|---|---|
| Primary somatosensory cortex: trunk region | 0,47244509 | 0,02681866 | 0,47677322 | 0,02416501 | 0,9 | 0,6508 | 0,8341 |
| anterior commissure: pars posterior | 0,52441578 | 0,02050409 | 0,52867327 | 0,02566559 | 0,8 | 0,5877 | 0,8165 |
| Primary visual cortex | 2,10831419 | 0,13940324 | 2,12528553 | 0,1080655 | 0,8 | 0,7268 | 0,8709 |
| Secondary visual cortex: mediomedial area | 1,68884951 | 0,07733081 | 1,70236726 | 0,09013005 | 0,8 | 0,6416 | 0,8341 |
| Claustrum | 0,30536355 | 0,01180902 | 0,30778612 | 0,01399021 | 0,8 | 0,5868 | 0,8165 |
| Claustrum: ventral part | 0,50504908 | 0,01421057 | 0,50904263 | 0,02267479 | 0,8 | 0,4986 | 0,7720 |
| flocculus (FL) | 0,83835293 | 0,05805165 | 0,84492247 | 0,04841709 | 0,8 | 0,7476 | 0,8835 |
| dentate nucleus | 0,32226255 | 0,01904787 | 0,32476299 | 0,01993666 | 0,8 | 0,7198 | 0,8709 |
| Ventral tenia tecta | 0,12596023 | 0,0078955 | 0,12680222 | 0,00821958 | 0,7 | 0,7703 | 0,9045 |
| facial nerve (cranial nerve 7) | 0,23702859 | 0,01014094 | 0,23859392 | 0,01411217 | 0,7 | 0,6955 | 0,8611 |
| Intermediate nucleus of the endopiriform claustrum | 0,64417452 | 0,0210887 | 0,64841744 | 0,0325974 | 0,7 | 0,6231 | 0,8278 |
| Primary somatosensory cortex: dysgranular zone | 0,3252095 | 0,01844976 | 0,3272734 | 0,01344181 | 0,6 | 0,7463 | 0,8835 |
| fasciculus retroflexus | 0,28206984 | 0,02221457 | 0,28357564 | 0,02244757 | 0,5 | 0,8518 | 0,9382 |
| Cingulate cortex: area 30 | 2,68478305 | 0,10327055 | 2,69819037 | 0,11702057 | 0,5 | 0,7274 | 0,8709 |
| Primary visual cortex: monocular area | 1,81632948 | 0,08311517 | 1,82488163 | 0,09439588 | 0,5 | 0,7823 | 0,9126 |
| Lateral orbital cortex | 3,34821435 | 0,11075199 | 3,36337554 | 0,15022961 | 0,5 | 0,7263 | 0,8709 |
| amygdala | 10,1273932 | 0,22564346 | 10,1710824 | 0,4075519 | 0,4 | 0,6570 | 0,8362 |
| Primary visual cortex: binocular area | 1,95057734 | 0,08703021 | 1,95730527 | 0,087445 | 0,3 | 0,8311 | 0,9382 |
| optic tract | 1,95938863 | 0,07390157 | 1,96586034 | 0,09614222 | 0,3 | 0,8204 | 0,9332 |
| Ventral nucleus of the endopiriform claustrum | 0,53910768 | 0,01407692 | 0,54059338 | 0,02295718 | 0,3 | 0,8002 | 0,9276 |
| simple lobule (lobule 6) | 4,90205705 | 0,27230306 | 4,91146571 | 0,24682192 | 0,2 | 0,9226 | 0,9623 |
| Cingulate cortex: area 29b | 0,40554654 | 0,02886456 | 0,40626403 | 0,02341987 | 0,2 | 0,9433 | 0,9623 |
| Parietal cortex: posterior area: rostral part | 0,10143006 | 0,00703225 | 0,10160605 | 0,00610671 | 0,2 | 0,9435 | 0,9623 |
| trunk of simple and crus 1 white matter | 0,16807181 | 0,00828012 | 0,1683396 | 0,00880986 | 0,2 | 0,9297 | 0,9623 |
| Dorsolateral entorhinal cortex | 2,77983199 | 0,17814586 | 2,78350983 | 0,14913899 | 0,1 | 0,9531 | 0,9637 |
| fastigial nucleus | 0,45647772 | 0,02178573 | 0,45701662 | 0,02300385 | 0,1 | 0,9461 | 0,9623 |
| Primary somatosensory cortex: barrel field | 8,6187193 | 0,43808676 | 8,62686895 | 0,42013671 | 0,1 | 0,9587 | 0,9640 |
| lobule 3: central lobule (dorsal) | 1,97733676 | 0,14051983 | 1,97585629 | 0,13042 | -0,1 | 0,9764 | 0,9764 |
| anterior lobule (lobules 4-5) | 1,58459671 | 0,08453176 | 1,58225587 | 0,07249397 | -0,1 | 0,9374 | 0,9623 |
| trunk of arbor vita | 4,69676415 | 0,15367346 | 4,68789636 | 0,19397772 | -0,2 | 0,8800 | 0,9427 |
| Primary somatosensory cortex: shoulder region | 0,17711934 | 0,00999204 | 0,17676997 | 0,00952735 | -0,2 | 0,9223 | 0,9623 |
| inferior olivary complex | 0,41585699 | 0,03499256 | 0,41499709 | 0,03703876 | -0,2 | 0,9465 | 0,9623 |
| Dorsal intermediate entorhinal cortex | 1,77324248 | 0,11862926 | 1,76826135 | 0,08586394 | -0,3 | 0,9031 | 0,9612 |
| mammillary bodies | 0,60755459 | 0,03070072 | 0,60577129 | 0,04020818 | -0,3 | 0,8805 | 0,9427 |
| Primary somatosensory cortex: hindlimb region | 2,43049692 | 0,11528897 | 2,42335117 | 0,1317159 | -0,3 | 0,8679 | 0,9427 |

| | | | | | | | |
|---|---|---|---|---|---|---|---|
| Primary somatosensory cortex: forelimb region | 3,675154 | 0,20956433 | 3,6617634 | 0,17577627 | -0,4 | 0,8557 | 0,9382 |
| simple lobule white matter | 0,31767488 | 0,02122722 | 0,31626691 | 0,02093303 | -0,4 | 0,8542 | 0,9382 |
| copula: pyramis (lobule 8) | 2,49490175 | 0,19239295 | 2,48199775 | 0,15872447 | -0,5 | 0,8483 | 0,9382 |
| lateral ventricle | 3,27352525 | 0,4113628 | 3,25023415 | 0,37304463 | -0,7 | 0,8735 | 0,9427 |
| medulla | 29,8664483 | 1,15773263 | 29,6446688 | 1,50145477 | -0,7 | 0,6202 | 0,8278 |
| lobule 9: uvula | 2,9066195 | 0,25784125 | 2,88434053 | 0,22632833 | -0,8 | 0,8072 | 0,9298 |
| anterior lobule white matter | 0,04969825 | 0,00274889 | 0,04931223 | 0,00257032 | -0,8 | 0,6952 | 0,8611 |
| lobule 9 white matter | 0,28003849 | 0,02923447 | 0,27767695 | 0,02421647 | -0,8 | 0,8180 | 0,9332 |
| cerebellar peduncle: middle | 1,36584665 | 0,05171403 | 1,35355675 | 0,06396173 | -0,9 | 0,5341 | 0,7866 |
| lobule 10 white matter | 0,07252789 | 0,0036706 | 0,07181939 | 0,00628085 | -1,0 | 0,6505 | 0,8341 |
| Piriform cortex | 10,3769459 | 0,34110397 | 10,2631986 | 0,5897951 | -1,1 | 0,4376 | 0,7413 |
| trunk of lobules 1-3 white matter | 0,15022916 | 0,00631088 | 0,14848571 | 0,01094243 | -1,2 | 0,5199 | 0,7819 |
| crus 1: ansiform lobule (lobule 6) | 4,52595474 | 0,23727716 | 4,46960984 | 0,2321898 | -1,2 | 0,5132 | 0,7783 |
| olfactory tubercle | 4,17164009 | 0,22181799 | 4,11753327 | 0,33641659 | -1,3 | 0,5491 | 0,7866 |
| cerebellar peduncle: inferior | 0,80910577 | 0,02151627 | 0,79839197 | 0,03955529 | -1,3 | 0,2621 | 0,6173 |
| lobule 3 white matter | 0,16277003 | 0,0132564 | 0,16059318 | 0,0115367 | -1,3 | 0,6434 | 0,8341 |
| Posterolateral cortical amygdaloid area | 0,92867969 | 0,02101157 | 0,91590342 | 0,04672273 | -1,4 | 0,2136 | 0,5541 |
| Rostral amygdalopiriform area | 0,43032013 | 0,02087232 | 0,42424883 | 0,02332603 | -1,4 | 0,4369 | 0,7413 |
| copula white matter | 0,06373694 | 0,00520305 | 0,06280716 | 0,00441079 | -1,5 | 0,6133 | 0,8278 |
| cuneate nucleus | 0,2895919 | 0,02340087 | 0,28518825 | 0,02217196 | -1,5 | 0,6012 | 0,8278 |
| lobules 6-7 white matter | 0,78639878 | 0,03433511 | 0,77333321 | 0,04634698 | -1,7 | 0,3352 | 0,6882 |
| crus 1 white matter | 0,29016472 | 0,01947232 | 0,2845222 | 0,02092631 | -1,9 | 0,4348 | 0,7413 |
| medial lemniscus/medial longitudinal fasciculus | 2,8271953 | 0,15968473 | 2,76912384 | 0,16166873 | -2,1 | 0,3245 | 0,6788 |
| lobules 1-2: lingula and central lobule (ventral) | 1,60084766 | 0,13568707 | 1,5651507 | 0,11193986 | -2,2 | 0,4579 | 0,7447 |
| lobule 6: declive | 2,83152436 | 0,11501466 | 2,76803218 | 0,17744593 | -2,2 | 0,1850 | 0,5181 |
| superior olivary complex | 0,74849153 | 0,04079824 | 0,73141862 | 0,03535841 | -2,3 | 0,2485 | 0,6111 |
| habenular commissure | 0,02846981 | 0,00380206 | 0,02777206 | 0,00411348 | -2,5 | 0,6192 | 0,8278 |
| paramedian lobule (lobule 7) | 4,27320435 | 0,40469988 | 4,16460621 | 0,27443275 | -2,5 | 0,4394 | 0,7413 |
| pontine nucleus | 0,99000204 | 0,04672016 | 0,9620139 | 0,05132905 | -2,8 | 0,1193 | 0,4097 |
| crus 2: ansiform lobule (lobule 7) | 4,32468159 | 0,39521802 | 4,19998401 | 0,27800933 | -2,9 | 0,3674 | 0,6952 |
| trunk of crus 2 and paramedian white matter | 0,40788171 | 0,03199498 | 0,39477231 | 0,02722624 | -3,2 | 0,2569 | 0,6173 |
| Cortex-amygdala transition zones | 0,84117256 | 0,03628263 | 0,81327345 | 0,0599173 | -3,3 | 0,0761 | 0,3070 |
| lobule 8: pyramis | 1,61629538 | 0,0857099 | 1,55546581 | 0,10761284 | -3,8 | 0,0761 | 0,3070 |
| crus 2 white matter | 0,21907404 | 0,02183394 | 0,21040216 | 0,01644418 | -4,0 | 0,2644 | 0,6173 |
| paramedian lobule | 0,11406017 | 0,01122603 | 0,1090758 | 0,00863994 | -4,4 | 0,2161 | 0,5541 |
| lobule 7: tuber (or folium) | 1,15129245 | 0,10502856 | 1,09839546 | 0,0992639 | -4,6 | 0,1745 | 0,5013 |
| lobule 1-2 white matter | 0,05662321 | 0,00603894 | 0,05400888 | 0,00613429 | -4,6 | 0,2444 | 0,6093 |
| corticospinal tract/pyramids | 2,12628059 | 0,17515514 | 2,01220044 | 0,13987493 | -5,4 | 0,0805 | 0,3118 |

### Supplemental Table S2

| Relative volumes (% of total brain) | STD | | ENR | | | |
|---|---|---|---|---|---|---|
| Atlas region of interest | Mean | SD | Mean | SD | Effect size | p-value |
| CA1Or | 0,44738035 | 0,00910027 | 0,47398411 | 0,01505453 | 2,92 | 2,9915E-07 |
| CA1Py | 0,22441587 | 0,00643615 | 0,23774594 | 0,00827928 | 2,07 | 3,6087E-05 |
| SLu | 0,13878779 | 0,00286007 | 0,14345238 | 0,00336634 | 1,63 | 0,0004 |
| Cingulate cortex: area 25 | 0,12265596 | 0,00561376 | 0,13176316 | 0,00854244 | 1,62 | 0,0006 |
| CA3Rad | 0,41460029 | 0,01145177 | 0,4331594 | 0,00906609 | 1,62 | 0,0005 |
| CA1Rad | 0,54790077 | 0,01898942 | 0,57769198 | 0,01798671 | 1,57 | 0,0006 |
| anterior commissure: pars anterior | 0,25261468 | 0,00687237 | 0,26293313 | 0,0075172 | 1,50 | 0,0008 |
| CA2Py | 0,04037863 | 0,00147265 | 0,04257741 | 0,00122028 | 1,49 | 0,0009 |
| CA2Rad | 0,11028174 | 0,0040401 | 0,11624448 | 0,00426438 | 1,48 | 0,0010 |
| subependymale zone / rhinocele | 0,01198026 | 0,00058852 | 0,01282122 | 0,00058385 | 1,43 | 0,0012 |
| LMol | 0,52288186 | 0,01735023 | 0,54681254 | 0,01745368 | 1,38 | 0,0016 |
| CA3Py Outer | 0,20913743 | 0,00377114 | 0,21428841 | 0,00569167 | 1,37 | 0,0026 |
| CA2Or | 0,10383407 | 0,004088 | 0,10936231 | 0,00304886 | 1,35 | 0,0018 |
| Medial orbital cortex | 0,40766066 | 0,01553291 | 0,42686664 | 0,01970585 | 1,24 | 0,0043 |
| CA3Py Inner | 0,02509506 | 0,00104391 | 0,02636898 | 0,00070166 | 1,22 | 0,0037 |
| trunk of lobules 6-8 white matter | 0,02970808 | 0,00139949 | 0,02751998 | 0,00194853 | -1,56 | 0,0007 |
| lateral septum | 0,66098515 | 0,01358793 | 0,67665128 | 0,02156351 | 1,15 | 0,0097 |
| Dorsal tenia tecta | 0,22792565 | 0,01115008 | 0,24047284 | 0,01628987 | 1,13 | 0,0098 |
| subiculum | 0,7622483 | 0,02393725 | 0,78901495 | 0,01822765 | 1,12 | 0,0065 |
| GrDG | 0,27124138 | 0,01229777 | 0,28366496 | 0,00707545 | 1,01 | 0,0112 |
| MoDG | 0,94283215 | 0,03541338 | 0,97565599 | 0,02540149 | 0,93 | 0,0181 |
| PoDG | 0,11436346 | 0,00609634 | 0,11995862 | 0,00334811 | 0,92 | 0,0182 |
| crus 1: ansiform lobule (lobule 6) | 1,00461237 | 0,03391635 | 0,97886771 | 0,03197685 | -0,76 | 0,0492 |
| Cortex-amygdala transition zones | 0,1869921 | 0,01097557 | 0,17823375 | 0,01301665 | -0,80 | 0,0463 |
| corticospinal tract/pyramids | 0,47224385 | 0,03849465 | 0,44085124 | 0,02740268 | -0,82 | 0,0330 |
| lobule 8 white matter | 0,02359414 | 0,00162314 | 0,02199149 | 0,00152367 | -0,99 | 0,0142 |
| cerebellar peduncle: middle | 0,3032688 | 0,00653291 | 0,2965577 | 0,0110886 | -1,03 | 0,0218 |
| cerebellar peduncle: inferior | 0,17972499 | 0,00450286 | 0,1748798 | 0,00582722 | -1,08 | 0,0111 |
| lobule 8: pyramis | 0,35892688 | 0,01688104 | 0,3407083 | 0,02005376 | -1,08 | 0,0100 |
| trunk of arbor vita | 1,04290461 | 0,01382358 | 1,02694431 | 0,02458179 | -1,15 | 0,0121 |
| pontine nucleus | 0,21980943 | 0,00767796 | 0,2107907 | 0,009877 | -1,17 | 0,0063 |
| corpus callosum | 2,82293684 | 0,04442591 | 2,85810932 | 0,08102896 | 0,79 | 0,0787 |
| fimbria | 0,69352628 | 0,01984717 | 0,70922122 | 0,03179942 | 0,79 | 0,0659 |
| bed nucleus of stria terminalis | 0,27996198 | 0,00603217 | 0,28447721 | 0,00849073 | 0,75 | 0,0697 |
| Accessory olfactory bulb: granule cell layer | 0,04111303 | 0,00219526 | 0,04270806 | 0,0018849 | 0,73 | 0,0560 |
| Anterior olfactory nucleus | 0,49792298 | 0,01321634 | 0,50733119 | 0,01507494 | 0,71 | 0,0701 |
| striatum | 4,44427833 | 0,05809884 | 4,4853862 | 0,08424031 | 0,71 | 0,0876 |

| | | | | | | |
|---|---|---|---|---|---|---|
| nucleus accumbens | 0,94750149 | 0,03505151 | 0,96973695 | 0,02675862 | 0,63 | 0,0866 |
| Secondary somatosensory cortex | 1,31655587 | 0,03761262 | 1,33966321 | 0,0539378 | 0,61 | 0,1333 |
| olfactory peduncle | 0,56440253 | 0,01705192 | 0,57471186 | 0,01495285 | 0,60 | 0,1050 |
| Cingulum | 0,19571438 | 0,00750644 | 0,20023389 | 0,00776932 | 0,60 | 0,1140 |
| Cingulate cortex: area 32 | 0,5645866 | 0,03631371 | 0,58620397 | 0,02520748 | 0,60 | 0,1029 |
| Temporal association area | 0,56698475 | 0,01527465 | 0,57596102 | 0,01797444 | 0,59 | 0,1309 |
| Olfactory bulb: mitral cell layer | 0,22640847 | 0,00676993 | 0,23014878 | 0,01067461 | 0,55 | 0,1879 |
| Olfactory bulb: granule cell layer | 0,99268354 | 0,03553033 | 1,0123 | 0,04286836 | 0,55 | 0,1564 |
| Frontal cortex: area 3 | 0,15535296 | 0,00387593 | 0,15744985 | 0,00515953 | 0,54 | 0,1744 |
| CA30r | 0,66771565 | 0,01666502 | 0,67654859 | 0,02127021 | 0,53 | 0,1782 |
| Olfactory bulb: internal plexiform layer | 0,16199689 | 0,00472587 | 0,16445059 | 0,00769339 | 0,52 | 0,2201 |
| Olfactory bulb: external plexiform layer | 1,5640047 | 0,04194853 | 1,58439622 | 0,06757904 | 0,49 | 0,2482 |
| internal capsule | 0,70763511 | 0,01517865 | 0,71494952 | 0,02424966 | 0,48 | 0,2508 |
| Ectorhinal cortex | 0,58495642 | 0,01784488 | 0,59330285 | 0,01889297 | 0,47 | 0,2136 |
| Cingulate cortex: area 24a | 0,4221665 | 0,01874869 | 0,43062037 | 0,02172767 | 0,45 | 0,2381 |
| Frontal association cortex | 1,60304255 | 0,04324518 | 1,62212261 | 0,05813322 | 0,44 | 0,2653 |
| Medial preoptic nucleus | 0,03825303 | 0,00199315 | 0,0390947 | 0,00226224 | 0,42 | 0,2658 |
| Cingulate cortex: area 29c | 0,44133804 | 0,01429861 | 0,44694439 | 0,01769505 | 0,39 | 0,3098 |
| Primary somatosensory cortex | 0,90605089 | 0,01631798 | 0,91238973 | 0,03206603 | 0,39 | 0,3940 |
| Ventral orbital cortex | 0,35150431 | 0,01017748 | 0,35541149 | 0,01220808 | 0,38 | 0,3161 |
| thalamus | 3,87434828 | 0,15734938 | 3,93259137 | 0,14168544 | 0,37 | 0,3071 |
| fundus of striatum | 0,03838909 | 0,00090856 | 0,03871959 | 0,0015158 | 0,36 | 0,3908 |
| Primary somatosensory cortex: upper lip region | 1,15167414 | 0,02305773 | 1,1597968 | 0,04848427 | 0,35 | 0,4544 |
| Cingulate cortex: area 24b | 0,37979944 | 0,02061472 | 0,3866977 | 0,0187717 | 0,33 | 0,3549 |
| Dorsolateral orbital cortex | 0,20575551 | 0,00781359 | 0,20819033 | 0,00971712 | 0,31 | 0,4175 |
| mammilothalamic tract | 0,05516709 | 0,00160321 | 0,05564772 | 0,00241413 | 0,30 | 0,4614 |
| Olfactory bulb: glomerular layer | 1,03584222 | 0,02490166 | 1,04328895 | 0,04555959 | 0,30 | 0,4964 |
| fornix | 0,16423679 | 0,00272292 | 0,16504416 | 0,00392453 | 0,30 | 0,4596 |
| Secondary auditory cortex: ventral area | 0,32665246 | 0,01015156 | 0,32929212 | 0,01157154 | 0,26 | 0,4882 |
| Lateral parietal association cortex | 0,04616392 | 0,00168055 | 0,0465768 | 0,00220108 | 0,25 | 0,5271 |
| pre-para subiculum | 0,48304598 | 0,01674434 | 0,48692561 | 0,01922506 | 0,23 | 0,5368 |
| cerebral aqueduct | 0,08306235 | 0,00699859 | 0,08456231 | 0,01229647 | 0,21 | 0,6190 |
| posterior commissure | 0,03379574 | 0,00257577 | 0,03428892 | 0,00245582 | 0,19 | 0,5953 |
| globus pallidus | 0,73565098 | 0,03449633 | 0,7415363 | 0,02799913 | 0,17 | 0,6272 |
| Perirhinal cortex | 0,53605704 | 0,02122387 | 0,53950798 | 0,02036175 | 0,16 | 0,6518 |
| cerebellar peduncle: superior | 0,2304521 | 0,0110735 | 0,23224493 | 0,01034429 | 0,16 | 0,6517 |
| medial septum | 0,312558 | 0,00739103 | 0,31374429 | 0,00666303 | 0,16 | 0,6526 |
| Secondary motor cortex | 1,36591413 | 0,03433267 | 1,37131632 | 0,04383035 | 0,16 | 0,6823 |
| Accessory olfactory bulb: glomerular, external plexiform and mitral cell layer | 0,08244534 | 0,00533588 | 0,08328233 | 0,0048568 | 0,16 | 0,6604 |

| | | | | | | |
|---|---|---|---|---|---|---|
| paraflocculus white matter | 0,04014934 | 0,00325755 | 0,04065042 | 0,00333664 | 0,15 | 0,6731 |
| periaqueductal grey | 0,85166742 | 0,04911357 | 0,85842582 | 0,04127725 | 0,14 | 0,6962 |
| Cingulate cortex: area 24a' | 0,19096889 | 0,01080313 | 0,19237991 | 0,00872847 | 0,13 | 0,7093 |
| midbrain | 3,14348497 | 0,12350305 | 3,15865364 | 0,12916186 | 0,12 | 0,7370 |
| fourth ventricle | 0,16193909 | 0,01271838 | 0,1634275 | 0,0123196 | 0,12 | 0,7453 |
| stria medullaris | 0,16400581 | 0,00696224 | 0,16478914 | 0,0057345 | 0,11 | 0,7486 |
| Medial parietal association cortex | 0,10056525 | 0,00332668 | 0,10089062 | 0,00455918 | 0,10 | 0,8030 |
| lobules 4-5 white matter | 0,11189182 | 0,00966397 | 0,11278515 | 0,00718466 | 0,09 | 0,7895 |
| colliculus: superior | 1,9736193 | 0,08772362 | 1,98156468 | 0,07648035 | 0,09 | 0,7979 |
| cerebral peduncle | 0,54327222 | 0,01452435 | 0,54452739 | 0,01408934 | 0,09 | 0,8103 |
| lobule 10: nodulus | 0,26778938 | 0,00958608 | 0,26856496 | 0,01521442 | 0,08 | 0,8445 |
| flocculus white matter | 0,01081835 | 0,00099462 | 0,01087919 | 0,00067908 | 0,06 | 0,8584 |
| interpeduncular nucleus | 0,05938421 | 0,00376693 | 0,05957923 | 0,00407513 | 0,05 | 0,8881 |
| Caudomedial entorhinal cortex | 1,09763694 | 0,06079622 | 1,10056413 | 0,04141552 | 0,05 | 0,8882 |
| lateral olfactory tract | 0,34228533 | 0,01009933 | 0,34271957 | 0,01444274 | 0,04 | 0,9138 |
| Medial entorhinal cortex | 0,14983239 | 0,0091323 | 0,15010216 | 0,0059292 | 0,03 | 0,9310 |
| basal forebrain | 1,24366424 | 0,03339021 | 1,24463325 | 0,03409634 | 0,03 | 0,9363 |
| hypothalamus | 2,42264075 | 0,02705832 | 2,42309334 | 0,05918566 | 0,02 | 0,9721 |
| Primary somatosensory cortex: jaw region | 0,14618361 | 0,00484428 | 0,1462619 | 0,00540177 | 0,02 | 0,9652 |
| Insular region: not subdivided | 1,46394603 | 0,02808505 | 1,46396816 | 0,04852861 | 0,00 | 0,9985 |
| nucleus interpositus | 0,09682453 | 0,00384371 | 0,09677416 | 0,00381049 | -0,01 | 0,9711 |
| paraflocculus (PFL) | 0,48327509 | 0,06426954 | 0,48216278 | 0,07215322 | -0,02 | 0,9628 |
| Cingulate cortex: area 24b' | 0,13201409 | 0,004965 | 0,13192343 | 0,00546237 | -0,02 | 0,9605 |
| Secondary visual cortex: lateral area | 0,5458248 | 0,01507436 | 0,54550508 | 0,01721273 | -0,02 | 0,9546 |
| Claustrum: dorsal part | 0,04852304 | 0,00152619 | 0,04848035 | 0,00178128 | -0,03 | 0,9404 |
| colliculus: inferior | 1,29234509 | 0,04869206 | 1,29073942 | 0,04463793 | -0,03 | 0,9262 |
| lobules 4-5: culmen (ventral and dorsal) | 0,96114964 | 0,04091632 | 0,95971624 | 0,04511902 | -0,04 | 0,9244 |
| Primary motor cortex | 1,59645858 | 0,06176136 | 1,59421956 | 0,05643482 | -0,04 | 0,9189 |
| ventral tegmental decussation | 0,02874792 | 0,0021236 | 0,02865576 | 0,00198772 | -0,04 | 0,9034 |
| Cingulate cortex: area 29a | 0,15867784 | 0,00537937 | 0,15841997 | 0,0062009 | -0,05 | 0,8978 |
| Ventral intermediate entorhinal cortex | 0,22633802 | 0,0145385 | 0,22560927 | 0,01014785 | -0,05 | 0,8840 |
| Amygdalopiriform transition area | 0,22704863 | 0,01416639 | 0,22626567 | 0,01112935 | -0,06 | 0,8739 |
| third ventricle | 0,25902556 | 0,00796296 | 0,25855735 | 0,01289557 | -0,06 | 0,8875 |
| Secondary visual cortex: mediolateral area | 0,22442467 | 0,00844987 | 0,22388926 | 0,0104841 | -0,06 | 0,8678 |
| Posteromedial cortical amygdaloid area | 0,29379332 | 0,01643487 | 0,2924533 | 0,01363229 | -0,08 | 0,8163 |
| Medial amygdala | 0,29415769 | 0,00645833 | 0,29360269 | 0,01119274 | -0,09 | 0,8407 |
| Primary visual cortex | 0,46795955 | 0,02461129 | 0,46570307 | 0,02098919 | -0,09 | 0,7948 |
| Secondary auditory cortex: dorsal area | 0,28927485 | 0,0069648 | 0,2886341 | 0,00825903 | -0,09 | 0,8068 |
| flocculus (FL) | 0,1861171 | 0,01073963 | 0,18506852 | 0,00811852 | -0,10 | 0,7784 |

| | | | | | | |
|---|---|---|---|---|---|---|
| Primary somatosensory cortex: trunk region | 0,10486817 | 0,00412282 | 0,10445863 | 0,00441446 | -0,10 | 0,7868 |
| Ventral tenia tecta | 0,02799315 | 0,00196156 | 0,02778481 | 0,00167338 | -0,11 | 0,7633 |
| fasciculus retroflexus | 0,06262173 | 0,00433847 | 0,06208884 | 0,00403431 | -0,12 | 0,7314 |
| dentate nucleus | 0,0715278 | 0,00292835 | 0,07111283 | 0,00298764 | -0,14 | 0,6972 |
| pons | 3,94899239 | 0,11271006 | 3,93213594 | 0,12508215 | -0,15 | 0,6867 |
| Primary somatosensory cortex: dysgranular zone | 0,07219151 | 0,0029397 | 0,07171079 | 0,00231319 | -0,16 | 0,6401 |
| stria terminalis | 0,20702887 | 0,00297221 | 0,20652494 | 0,00611512 | -0,17 | 0,7148 |
| lateral ventricle | 0,72708839 | 0,09079196 | 0,71132638 | 0,06990017 | -0,17 | 0,6189 |
| Primary auditory cortex | 0,29876823 | 0,00633721 | 0,29763714 | 0,00819396 | -0,18 | 0,6438 |
| anterior commissure: pars posterior | 0,11646334 | 0,0036081 | 0,11580678 | 0,00381773 | -0,18 | 0,6205 |
| Secondary visual cortex: mediomedial area | 0,37493824 | 0,01081891 | 0,3729592 | 0,015992 | -0,18 | 0,6497 |
| Dorsal nucleus of the endopiriform | 0,3349776 | 0,00788379 | 0,33351336 | 0,00857635 | -0,19 | 0,6155 |
| inferior olivary complex | 0,09235811 | 0,00763524 | 0,09092253 | 0,0076848 | -0,19 | 0,6054 |
| Intermediate nucleus of the endopiriform claustrum | 0,14312234 | 0,00551119 | 0,14207427 | 0,00615159 | -0,19 | 0,6092 |
| Dorsolateral entorhinal cortex | 0,61734376 | 0,03746476 | 0,60991583 | 0,0289215 | -0,20 | 0,5708 |
| Cingulate cortex: area 29b | 0,09000793 | 0,00513291 | 0,08899 | 0,00397401 | -0,20 | 0,5708 |
| Claustrum: ventral part | 0,11219316 | 0,0032716 | 0,11154288 | 0,00414521 | -0,20 | 0,6048 |
| facial nerve (cranial nerve 7) | 0,05263991 | 0,00185768 | 0,05225473 | 0,0023299 | -0,21 | 0,5884 |
| Parietal cortex: posterior area: rostral part | 0,02251013 | 0,00119345 | 0,0222574 | 0,00110909 | -0,21 | 0,5556 |
| lobule 9 white matter | 0,0621462 | 0,00584209 | 0,06082306 | 0,00486974 | -0,23 | 0,5222 |
| Claustrum | 0,06780397 | 0,00156091 | 0,06742481 | 0,00204563 | -0,24 | 0,5318 |
| Primary visual cortex: monocular area | 0,40330799 | 0,01390068 | 0,39985999 | 0,01801402 | -0,25 | 0,5219 |
| Dorsal intermediate entorhinal cortex | 0,39382157 | 0,02506886 | 0,38751628 | 0,01754221 | -0,25 | 0,4692 |
| lobule 3: central lobule (dorsal) | 0,43876982 | 0,02283475 | 0,43270136 | 0,02173787 | -0,27 | 0,4630 |
| copula: pyramis (lobule 8) | 0,55380597 | 0,03767121 | 0,54359864 | 0,02725056 | -0,27 | 0,4379 |
| lobule 9: uvula | 0,64492456 | 0,04713023 | 0,63179586 | 0,04431746 | -0,28 | 0,4413 |
| habenular commissure | 0,00632786 | 0,0008747 | 0,00608133 | 0,0008709 | -0,28 | 0,4405 |
| Ventral nucleus of the endopiriform claustrum | 0,11979343 | 0,00443465 | 0,1184701 | 0,00450212 | -0,30 | 0,4164 |
| trunk of simple and crus 1 white matter | 0,03732077 | 0,00146947 | 0,03687488 | 0,00141873 | -0,30 | 0,4046 |
| Primary visual cortex: binocular area | 0,43309898 | 0,01378487 | 0,42891229 | 0,0165336 | -0,30 | 0,4250 |
| simple lobule (lobule 6) | 1,08792283 | 0,0349197 | 1,07563716 | 0,03134965 | -0,35 | 0,3306 |
| fastigial nucleus | 0,10134663 | 0,00338382 | 0,1001289 | 0,00407834 | -0,36 | 0,3470 |
| cuneate nucleus | 0,06431543 | 0,00502328 | 0,06250335 | 0,00478181 | -0,36 | 0,3232 |
| copula white matter | 0,01415163 | 0,00107337 | 0,01376167 | 0,0009019 | -0,36 | 0,3113 |
| simple lobule white matter | 0,07049897 | 0,00338914 | 0,06924371 | 0,00317815 | -0,37 | 0,3098 |
| lobule 3 white matter | 0,03612693 | 0,00253026 | 0,03518295 | 0,00227096 | -0,37 | 0,3033 |
| Primary somatosensory cortex: shoulder region | 0,03931755 | 0,00157488 | 0,03872963 | 0,00178642 | -0,37 | 0,3230 |
| Primary somatosensory cortex: barrel field | 1,91317376 | 0,06099018 | 1,88988105 | 0,06685754 | -0,38 | 0,3089 |

| | | | | | | |
|---|---|---|---|---|---|---|
| Primary somatosensory cortex: forelimb region | 0,81587495 | 0,03543652 | 0,80221903 | 0,02932032 | -0,39 | 0,2832 |
| lobules 1-2: lingula and central lobule (ventral) | 0,35582059 | 0,03270795 | 0,34308125 | 0,02502624 | -0,39 | 0,2741 |
| paramedian lobule (lobule 7) | 0,94904995 | 0,09013858 | 0,9122896 | 0,05108316 | -0,41 | 0,2424 |
| amygdala | 2,24971 | 0,05139562 | 2,22834953 | 0,05805858 | -0,42 | 0,2726 |
| anterior lobule (lobules 4-5) | 0,35173763 | 0,0121701 | 0,34663803 | 0,01159302 | -0,42 | 0,2542 |
| Cingulate cortex: area 30 | 0,59609202 | 0,01178008 | 0,59112351 | 0,01778139 | -0,42 | 0,3035 |
| Primary somatosensory cortex: hindlimb region | 0,53967954 | 0,0198941 | 0,53086642 | 0,02293923 | -0,44 | 0,2455 |
| crus 2: ansiform lobule (lobule 7) | 0,96087458 | 0,09180226 | 0,91977654 | 0,04686594 | -0,45 | 0,2001 |
| olfactory tubercle | 0,92697851 | 0,05494312 | 0,90216453 | 0,07023297 | -0,45 | 0,2482 |
| anterior lobule white matter | 0,01103469 | 0,00050074 | 0,0108049 | 0,00049537 | -0,46 | 0,2166 |
| mammillary bodies | 0,13487469 | 0,00461294 | 0,13270163 | 0,00746167 | -0,47 | 0,2634 |
| optic tract | 0,43506082 | 0,00931431 | 0,43059483 | 0,01400658 | -0,48 | 0,2432 |
| Lateral orbital cortex | 0,74350304 | 0,01301516 | 0,7368674 | 0,02369613 | -0,51 | 0,2491 |
| lobule 1-2 white matter | 0,01258642 | 0,00141589 | 0,01184108 | 0,00137679 | -0,53 | 0,1588 |
| crus 2 white matter | 0,04864747 | 0,00478432 | 0,04608574 | 0,00316053 | -0,54 | 0,1367 |
| Rostral amygdalopiriform area | 0,09559943 | 0,00482547 | 0,09298044 | 0,00496822 | -0,54 | 0,1507 |
| Piriform cortex | 2,30614823 | 0,10368765 | 2,24906715 | 0,12183138 | -0,55 | 0,1552 |
| paramedian lobule | 0,02532766 | 0,00245177 | 0,02389341 | 0,00169878 | -0,58 | 0,1083 |
| trunk of crus 2 and paramedian white matter | 0,0905722 | 0,00676048 | 0,08649115 | 0,00537891 | -0,60 | 0,1022 |
| medulla | 6,63317162 | 0,21763601 | 6,49261775 | 0,20591375 | -0,65 | 0,0882 |
| superior olivary complex | 0,16627765 | 0,00918908 | 0,16026563 | 0,00661249 | -0,65 | 0,0770 |
| trunk of lobules 1-3 white matter | 0,03336585 | 0,00123522 | 0,03254565 | 0,00236787 | -0,66 | 0,1454 |
| crus 1 white matter | 0,06439129 | 0,0031403 | 0,0622969 | 0,00357585 | -0,67 | 0,0876 |
| lobules 6-7 white matter | 0,17469792 | 0,00782361 | 0,16940333 | 0,0082676 | -0,68 | 0,0798 |
| lobule 7: tuber (or folium) | 0,25557644 | 0,02162318 | 0,24062465 | 0,02015342 | -0,69 | 0,0695 |
| medial lemniscus/medial longitudinal fasciculus | 0,6277545 | 0,03016074 | 0,60659907 | 0,02827452 | -0,70 | 0,0662 |
| Posterolateral cortical amygdaloid area | 0,20638939 | 0,00799327 | 0,20070276 | 0,00926687 | -0,71 | 0,0710 |
| lobule 10 white matter | 0,01610024 | 0,00051307 | 0,01572066 | 0,00112197 | -0,74 | 0,1290 |
| lobule 6: declive | 0,62917626 | 0,0301765 | 0,60643893 | 0,0344151 | -0,75 | 0,0567 |

**Supplemental Table S3**

| Source ROI | Destination ROI |
|---|---|
| Accessory_olfactory_bulb:_granule_cell_layer | CA20r |
| Accessory_olfactory_bulb:_granule_cell_layer | Cingulate_cortex:_area_32 |
| Anterior_olfactory_nucleus | CA3Py_Outer |
| Anterior_olfactory_nucleus | SLu |
| CA10r | Olfactory_bulb:_granule_cell_layer |
| CA10r | Olfactory_peduncle |
| CA20r | Olfactory_bulb:_external_plexiform_layer_ |
| CA20r | Olfactory_bulb:_granule_cell_layer |
| CA20r | Olfactory_bulb:_internal_plexiform_layer_ |
| CA20r | Olfactory_bulb:_mitral_cell_layer |
| CA2Py | Olfactory_bulb:_granule_cell_layer |
| CA2Py | Olfactory_bulb:_internal_plexiform_layer_ |
| CA2Py | Olfactory_bulb:_mitral_cell_layer |
| CA2Rad | Olfactory_bulb:_granule_cell_layer |
| CA2Rad | Subiculum |
| CA3Py_Inner | Olfactory_bulb:_external_plexiform_layer_ |
| CA3Py_Inner | Olfactory_bulb:_granule_cell_layer |
| CA3Py_Inner | Olfactory_bulb:_internal_plexiform_layer_ |
| CA3Py_Inner | Olfactory_bulb:_mitral_cell_layer |
| CA3Py_Inner | Subiculum |
| CA3Py_Outer | Olfactory_peduncle |
| CA3Rad | Olfactory_bulb:_granule_cell_layer |
| CA3Rad | Subiculum |
| Claustrum | Olfactory_bulb:_external_plexiform_layer_ |
| Claustrum | Olfactory_bulb:_granule_cell_layer |
| Claustrum | Olfactory_bulb:_internal_plexiform_layer_ |
| Claustrum | Olfactory_bulb:_mitral_cell_layer |
| GrDG | Subiculum |
| LMol | Subiculum |
| MoDG | Subiculum |
| Olfactory_bulb:_external_plexiform_layer_ | Nucleus_accumbens |
| Olfactory_bulb:_granule_cell_layer | Bed_nucleus_of_stria_terminalis |
| Olfactory_bulb:_granule_cell_layer | Nucleus_accumbens |
| Olfactory_bulb:_granule_cell_layer | Subiculum |
| Olfactory_bulb:_internal_plexiform_layer_ | Nucleus_accumbens |
| Olfactory_bulb:_internal_plexiform_layer_ | Subiculum |
| Olfactory_bulb:_mitral_cell_layer | PoDG |
| Olfactory_bulb:_mitral_cell_layer | Bed_nucleus_of_stria_terminalis |
| Olfactory_bulb:_mitral_cell_layer | Nucleus_accumbens |
| Olfactory_bulb:_mitral_cell_layer | Subiculum |
| SLu | Olfactory_peduncle |
| Temporal_association_area | Anterior_commissure_pars_anterior |
| Anterior_commissure_pars_anterior | Corpus_callosum |
| Anterior_commissure_pars_anterior | Fimbria |
| Anterior_commissure_pars_anterior | Lateral_Septum |
| Fimbria | Olfactory_peduncle |
| Lateral_septum | Olfactory_peduncle |
